# Spread of pathological α-synuclein from urogenital nerves initiates multiple system atrophy-like symptoms

**DOI:** 10.1101/529594

**Authors:** Xuejing Wang, Mingming Ma, Erxi Wu, Dongyang Teng, Xin Yuan, Lebo Zhou, Rui Zhang, Jing Yao, Yongkang Chen, Danhao Xia, Robert K.F. Teng, Fengfei Wang, Jason H. Huang, Zhenyu Yue, Zhuohua Zhang, Junfang Teng, Beisha Tang, Xuebing Ding

## Abstract

Multiple system atrophy (MSA) is a fatal adult-onset movement disorder with autonomic failures, especially urogenital dysfunction. The neuropathological feature of MSA is the accumulation of misfolded α-synuclein (α-Syn) in the nervous system. Here, we show that misfolded α-Syn exist in nerve terminals in detrusor (DET) and external urethral sphincter (EUS) of patients with MSA. Moreover, α-Syn preformed fibrils inoculated into the EUS or DET in TgM83^+/−^ mice initiated the transmission of misfolded α-Syn from the lower urinary tract to brain, and these mice developed α-Syn inclusion pathology through micturition reflex pathways along with urinary dysfunction and motor impairments. These findings indicate that spreading of misfolded α-Syn from the autonomic control of the lower urinary tract to the brain via micturition reflex pathways induces autonomic failure and motor impairments. These results provide important new insights into the pathogenesis of MSA as well as highlight potential targets for early detection and therapeutics.

## Introduction

Multiple system atrophy (MSA) is a fatal, multisystem, neurodegenerative disorder characterized by a variable combination of rapidly progressive autonomic failures, ataxia, and parkinsonism. According to the most recent guidelines, autonomic failures featuring urogenital dysfunction, orthostatic hypotension, and respiratory disorder are premonitory symptoms and necessary for the diagnosis of MSA (Gilman et al., 2008). Retrospective data indicate that among autonomic failures, urological symptoms occur several years prior to the neurological symptoms in the majority of MSA patients (Beck, Betts, & Fowler, 1994; Jecmenica-Lukic, Poewe, Tolosa, & Wenning, 2012; Sakakibara et al., 2000). Urogenital dysfunction in patients with extrapyramidal symptoms is thought to help differentiate between Parkinson’s disease (PD) and MSA in early disease stages (Wenning et al., 1999). Consistently, neuropathological studies reveal that widespread pathological lesions of micturition reflex pathways, including the periaqueductal gray (PAG), Barrington’s nucleus (BN), intermediolateral columns (IML), Onuf’s nucleus of the spinal cord and so on, is present in central nervous system (CNS) of MSA patients (Stemberger, Poewe, Wenning, & Stefanova, 2010; VanderHorst et al., 2015). So far, few animal models of MSA have been established to display MSA-like urinary dysfunction and denervation-reinnervation of EAS simultaneously. It has been acknowledged that the cellular hallmark lesion of MSA is misfolded α-synuclein (α-Syn) accumulation within glial cytoplasmic inclusions along with neuronal inclusions (NIs) in central nervous system (CNS). Moreover, Watts et. al. reported that brain homogenates from MSA cases induced widespread deposits of phosphorylated α-Syn in the brains of MSA-inoculated mice, suggesting that α-Syn aggregates in the brains of MSA are transmissible (Watts et al., 2013).

In this study we show that misfolded α-Syn exist in nerve terminals in detrusor (DET) and external urethral sphincter (EUS) of patients with MSA. Also, we injected α-Syn preformed fibrils (PFFs) to the lower urinary tract of hemizygous TgM83^+/−^ mice, we observed the widespread α-Syn inclusion pathology from the autonomic control of the lower urinary tract to the brain along with urinary dysfunction and motor impairments

## Resultse

### Clinical characteristics of patients

Forty-five patients were diagnosed as MSA, PD, or PSP according to the consensus criteria (Gilman et al., 2008; Kalia & Lang, 2015; Litvan et al., 1996). Informed consent was obtained for each subject or their authorized surrogates on behalf of patients who lack decision-making ability. Clinical descriptions for each type of disease are summarized in Table 1. Among 32 patients (12 males and 20 females) with MSA, 13 patients were MSA with predominant parkinsonism (MSA-P) while 19 patients were MSA with predominant cerebellar ataxia (MSA-C). The mean ± SD of patients’ ages for these two types of MSA at the time of clinically diagnosed were 62.1 ± 7.1 y and 57.8 ± 6.6 y, respectively. The UMSARS scores were utilized for the evaluation of patients’ urological function. The score values (mean ± SD) of MSA-P and MSA-C were 44.4 ± 25.7 and 26.7 ± 15.3, respectively. In addition, these patients all had autonomic symptoms, including urological dysfunction, orthostatic dysregulation or chronic constipation (Low et al., 2015; Stefanova, Bucke, Duerr, & Wenning, 2009; Wenning et al., 2004). Furthermore, the positive rates of urodynamic examination and perianal electromyography in MSA were 88.1% and 81.0%, respectively. Altogether, these data indicate that urological dysfunction is specific and common in patients with MSA.

**Table 1.** Characteristics and the exam findings of patients

| Age | Sex | Diagnosis | Duration<br>(year) | MRI | UMSARS | | | | Urodynamic<br>examination† | Perianal<br>electromyography† | α-synuclein filament<br>in the bladder DET‡ | α-synuclein filament<br>in EUS§ |
| --- | --- | --- | --- | --- | --- | --- | --- | --- | --- | --- | --- | --- |
|  |  |  |  |  | UMSARS I | UMSARS II | UMSARS III | UMSARS IV |  |  |  |  |
| 54 | M | MSA-P | 1.5 | + | 2 | 5 | + | 1 | + | + | - | 0 |
| 57 | F | MSA-P | 8 | + | 32 | 50 | + | 4 | + | + | + | 0 |
| 63 | F | MSA-P | 2 | + | 14 | 18 | + | 2 | - | + | - | 0 |
| 59 | F | MSA-P | 4 | + | 21 | 18 | + | 3 | - | + | - | 0 |
| 46 | M | MSA-P | 2 | + | 9 | 14 | + | 2 | + | + | - | 0 |
| 68 | M | MSA-P | 7 | + | 20 | 31 | + | 3 | + | + | + | 0 |
| 66 | F | MSA-P | <1 | + | 7 | 12 | + | 1 | - | + | + | 0 |
| 70 | F | MSA-P | 3 | + | 32 | 46 | + | 4 | + | + | + | 0 |
| 66 | F | MSA-P | 3 | + | 18 | 24 | + | 3 | + | + | - | + |
| 72 | F | MSA-P | 5 | + | 34 | 42 | + | 4 | + | + | + | + |
| 63 | M | MSA-P | 2 | + | 7 | 13 | + | 2 | + | - | - | + |
| 64 | F | MSA-P | 7 | + | 33 | 42 | + | 5 | + | + | + | + |
| 59 | F | MSA-P | 4 | + | 18 | 15 | + | 4 | + | - | + | + |
| 59 | M | MSA-C | 2 | + | 10 | 13 | + | 2 | + | + | + | 0 |
| 50 | M | MSA-C | 1.5 | + | 2 | 3 | + | 1 | - | - | - | 0 |
| 47 | M | MSA-C | 8 | + | 30 | 21 | + | 4 | + | + | + | 0 |
| 63 | F | MSA-C | 2 | + | 21 | 14 | + | 1 | + | - | - | 0 |
| 50 | M | MSA-C | 2 | + | 5 | 6 | + | 1 | + | - | - | 0 |
| 61 | M | MSA-C | 3 | + | 20 | 18 | + | 4 | + | + | + | 0 |
| 61 | M | MSA-C | 5 | + | 8 | 9 | + | 2 | + | + | + | 0 |
| 51 | F | MSA-C | 3 | + | 8 | 6 | + | 1 | - | + | - | 0 |
| 58 | F | MSA-C | 3 | + | 4 | 5 | + | 2 | + | + | - | 0 |
| 50 | F | MSA-C | 2 | + | 12 | 14 | + | 2 | + | + | + | 0 |
| 64 | F | MSA-C | 2 | + | 14 | 24 | + | 4 | + | + | + | 0 |
| 62 | F | MSA-C | 3 | + | 14 | 18 | + | 3 | + | + | + | 0 |
| 64 | M | MSA-C | 3 | + | 10 | 11 | + | 2 | + | + | + | 0 |
| 64 | F | MSA-C | 3 | + | 30 | 12 | + | 4 | + | - | - | + |
| 52 | F | MSA-C | 8 | + | 33 | 22 | + | 5 | + | + | + | + |
| 53 | F | MSA-C* | 2 | + | 9 | 15 | + | 1 | + | + | - | + |
| 62 | F | MSA-C | 2 | + | 7 | 9 | + | 2 | + | - | + | + |
| 57 | F | MSA-C | 4 | + | 20 | 25 | + | 5 | + | - | + | + |
| 71 | M | MSA-C | 1 | + | 2 | 4 | + | 1 | + | + | + | - |
| 54 | F | PD | 10 | - | NA |  |  |  | - | + | - | 0 |
| 58 | F | PD | 3 | - | NA |  |  |  | - | - | - | 0 |
| 69 | F | PD | 10 | - | NA |  |  |  | + | - | - | 0 |
| 61 | M | PD | 6 | - | NA |  |  |  | - | - | - | 0 |
| 65 | F | PD | 10 | - | NA |  |  |  | - | - | - | 0 |
| 58 | F | PD | 4 | - | NA |  |  |  | - | - | - | - |
| 64 | M | PD | 8 | - | NA |  |  |  | - | + | - | - |
| 47 | M | PSP | 8 | + | NA |  |  |  | - | - | - | 0 |
| 69 | M | PSP | 3 | + | NA |  |  |  | - | - | - | 0 |
| 70 | M | PSP | 3 | + | NA |  |  |  | - | - | - | 0 |
| 69 | F | PSP | 3 | + | NA |  |  |  | + | - | - | 0 |
| 70 | F | PSP-C | 10 | + | NA |  |  |  | - | - | - | 0 |
| 67 | M | PSP | 4 | + | NA |  |  |  | - | - | - | - |

F, female; M, male; MSA-P, multiple system atrophy with predominant parkinsonism; MSA-C, multiple system atrophy with predominant cerebellar ataxia; PD, Parkinson's disease; PSP, progressive supranuclear palsy; PSP-C, progressive supranuclear palsy with predominant cerebellar ataxia; MRI, magnetic resonance imaging, including atrophy on MRI of putamen, middle cerebellar peduncle, pons, or cerebellum, -, absent; +, present; UMSARS, Unified Multiple System Atrophy Rating Scale (I. historical review of disease-related impairments; II. motor examination; III. autonomic examination, -, absent; +, present; IV. global disability scale); NA, not applicable.
\*possible MSA-C.
†-, normal findings; +, abnormal findings.
‡ DET, detrusor; -, absent; +, present.
§ EUS, external urethral sphincter; -, absent; +, present; 0: not examined.

### Detection of misfolded α-Syn in patients’ samples

We then investigated the deposits of misfolded α-Syn in MSA patients’ DET or EUS, using anti-α-Syn filament, anti-phosphorylated α-Syn (pα-Syn), and anti-aggregated-α-Syn (5G4) antibodies (Fig. 1A-H). Among 32 patients with MSA, 23 MSA cases exhibited deposits of misfolded α-Syn in biopsy tissues (Fig. 1A-Fand Table 1). Moreover, the portion of misfolded α-Syn in the triangle region, right wall, and left wall of bladder is similar (P > 0.05) and has no significant difference between MSA-P and MSA-C (P > 0.05). Among the patients examined, one had a urinary incontinence for 6 years before presence of movement deficits, and we made diagnosis for him as MSA after 8 months of movement deficits. A large amount of misfolded α-Syn was found in his DET and EUS. Most remarkably, no PD cases and PSP cases tested show misfolded α-Syn in bladder (Fig. 1G, H, I). Twenty control subjects didn’t show misfolded α-Syn in bladders either (Fig. 1I). Taken together, these results show that misfolded α-Syn exists in DET or EUS from 71.9% of the collected patients with MSA, while PD, PSP, and control subjects exhibit no detectable misfolded α-Syn in their bladders from this study.

**Figure 1.**
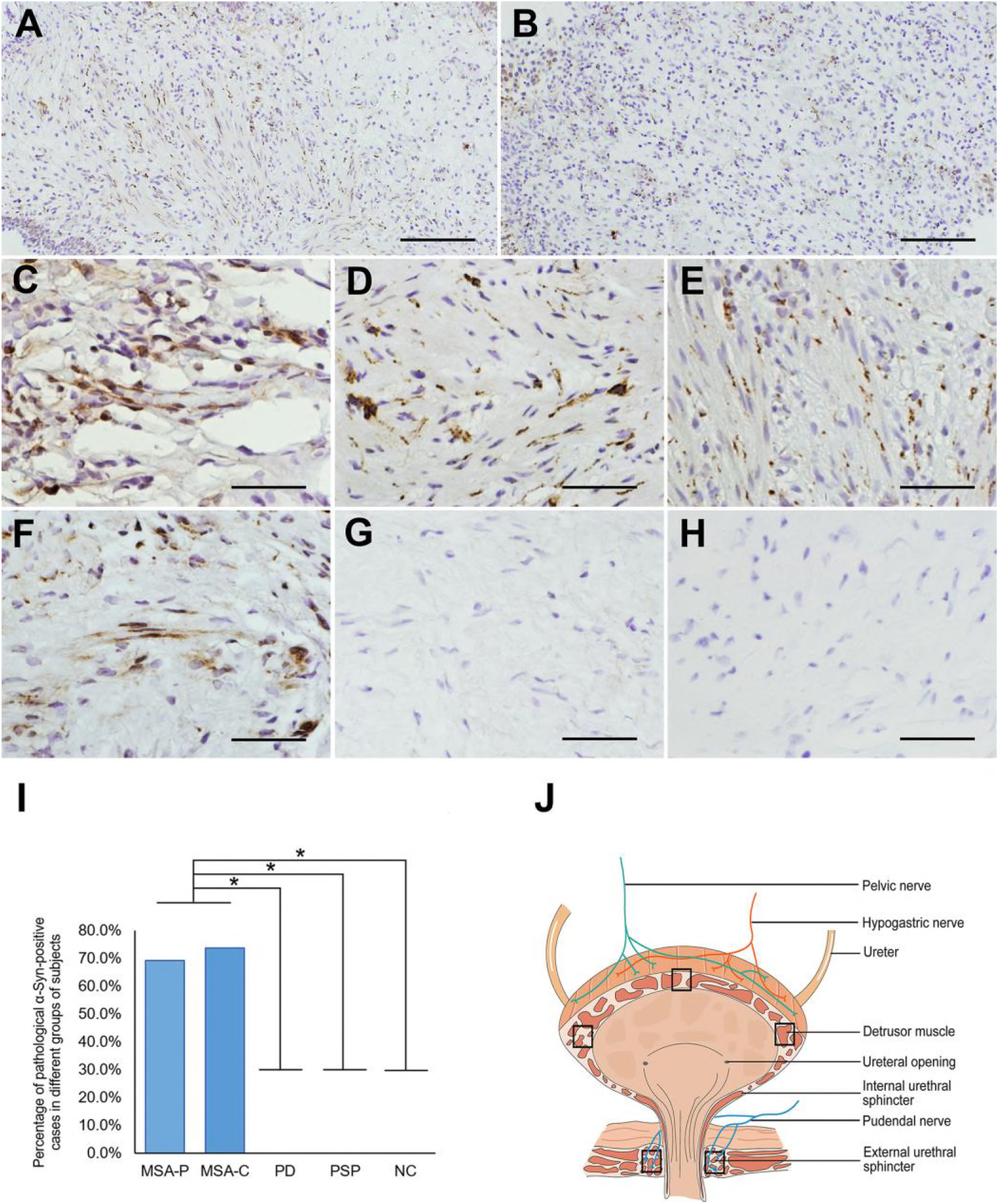
Immunohistochemical results of the sample tissues from subjects stained with anti-α-Syn filament antibody (MJFR14). (A-H) Representative images displayed misfolded α-Syn in DET of MSA-P (A, C-E) and MSA-C (B), and in EUS of MSA-P (F), but not in PSP (G) and PD (H). (C-E) Representative images displayed the right wall of MSA-P (C), the left wall of MSA-P (D) and the triangle region of MSA-P (E). (I) Histogram shows the percentage of cases who had α-Syn-positive inclusions in sample tissues in different groups. (J) Schematic displayed the anatomy of the lower urinary tract and sampling positions (DET and EUS). Statistical significance was analyzed employing the χ2 test. *P < 0.05. [Scale bar, 400 µm (A, B); 100 µm (C-H).]

### Identification of the micturition reflex pathways controlling EUS or DET using Fluoro-Gold (FG)

After we found that misfolded α-Syn proteins exist in DET and EUS of the detected patients, we then used Fluoro-Gold (FG) injection to trace the micturition reflex pathways controlling EUS or DET in mice. FG was injected into both sides of EUS or DET in TgM83^+/−^ and C57BL/6 mice; the sections from different parts of nervous system were detected at 14-day post-injection. FG-labeled neurons were detected in pelvic ganglia, spinal cord, pons, and midbrain bilaterally in both mouse models (Fig. 2). In spinal cord, we found that FG-labeled neurons also existed in T2 level, which has not been previously reported. At T2 level, the FG-labeled neurons mostly appeared in ventral horn and IML, which are closely associated with motor and autonomic functions. Among other levels of spinal cord, FG-labeled neurons gathered in the EUS motoneurons of lamina IX (ExU9) and sacral parasympathetic nucleus (SPSy) at S1 level, in EAS motoneurons of lamina IX (ExA9), ExU9, gluteal motoneurons of lamina IX (Gl9), lamina VII of the spinal gray (7Sp), and lateral spinal nucleus (LSp) at L6 level, in psoas motoneurons of lamina IX (Ps9), quadriceps motoneurons of lamina IX (Q9), intercalated nucleus (ICL), IML, and lumbar dorsal commissural nucleus (LDCom) at L2 level. In brain, FG-labeled neurons appeared in BN, PAG, and locus coeruleus (LC), and these nuclei have been reported to participate in micturition reflex pathways (Fowler, Griffiths, & de Groat, 2008). In addition, FG-labeled neurons were found to be in parvocellular reticular nucleus alpha (PCRtA), mesencephalic trigeminal nucleus (Me5), and red nucleus (RN), which are involved in the general locomotion, postural control, and modulation of certain sensory and autonomic functions. Taken together, these results suggest that the micturition reflex pathways controlling EUS and DET are connected not only with the autonomic nervous system but also with the central motor pathways.

**Figure 2.**
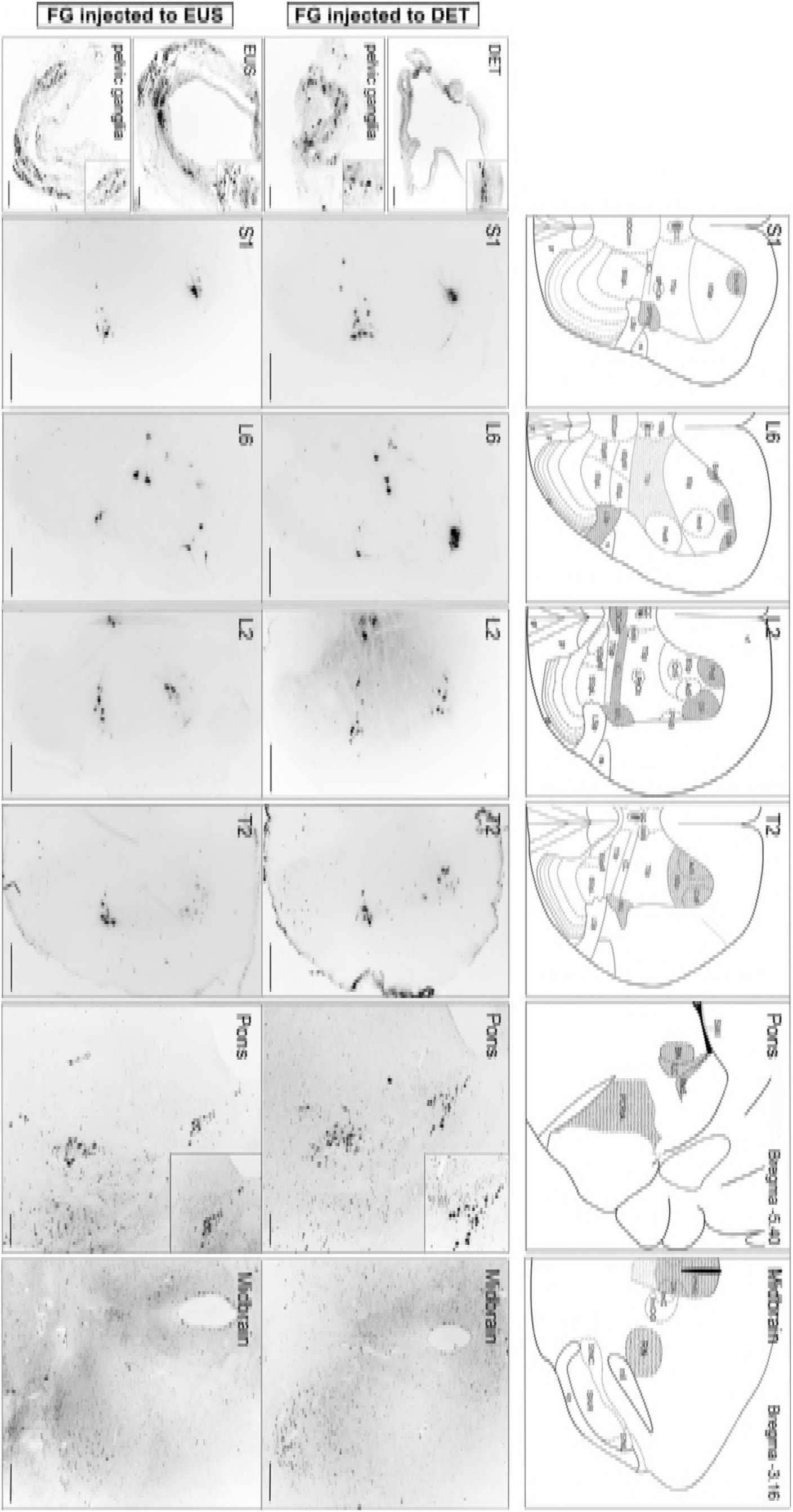
Representative images of FG-labeled neurons in DET-and EUS-FG C57BL/6 mice. All FG-labeled neurons appeared bilaterally while displayed one side. Schematics in the upper panel displayed the map of retrograde tracing areas (shaded areas) at different levels. Labeling appeared in DET-FG C57BL/6 mice (middle panel) and EUS-FG C57BL/6 mice (lower panel). Insets show a higher magnification relative to the main image. [Scale bars, 200 µm (DET, EUS, pons); 100 µm (pelvic ganglia); 500 µm (S1, L6, L2, T2, midbrain).] Abbreviation: 5SpL: lamina V of the spinal gray, lateral part; 5SpM: lamina V of the spinal gray, medial part; 6SpL: lamina VI of the spinal gray, lateral part; 6SpM: lamina VI of the spinal gray, medial part; 7Sp: lamina VII of the spinal gray; 8Sp: lamina VIII of the spinal gray; 10Sp: lamina X of the spinal gray; Ad9: adductor motoneurons of lamina IX Ax9: axial muscle motoneurons of lamina IX; BN: Barrington’s nucleus; CC: central canal; cp: cerebral peduncle, basal part; Cr9: cremaster motoneurons of lamina IX; csc: commissure of the superior colliculus; DET: detrusor; df: dorsal funiculus; Dk: nucleus of Darkschewitsch; dl: dorsolateral fasciculus (Lissauer); ExA9: external anal sphincter motoneurons of lamina IX; ExU9: external urethral sphincter motoneurons of lamina IX; EUS: external urethral sphincter; FG: Fluoro-Gold; Gl9: gluteal motoneurons of lamina IX; gr: gracile fasciculus; Hm9: hamstring motoneurons of lamina IX; ICL: intercalated nucleus; ICo9: intercostal muscle motoneurons of lamina IX; IML: intermediolateral columns; IMM: intermediomedial column; InCG: interstitial nucleus of Cajal, greater part; InC: interstitial nucleus of Cajal; LC: locus coeruleus; LDCom: lumbar dorsal commissural nucleus; LPrCb: lumbar precerebellar nucleus; LSp: lateral spinal nucleus; Me5: mesencephalic trigeminal nucleus; ml: medial lemniscus; PAG: periaqueductal gray; PCRtA: parvocellular reticular nucleus alpha; Pes9: pes motoneurons of lamina IX; Ps9: psoas motoneurons of lamina IX; Q9: quadriceps motoneurons of lamina IX; RN: red nucleus; rs: rubrospinal tract; SDCom: sacral dorsal commissural nucleus; SMV: superior medullary velum; SNC: substantia nigra pars compacta; SNL: substantia nigra, lateral part; SNR: substantia nigra, reticular part; SPrCb: sacral precerebellar nucleus; SPSy: sacral parasympathetic nucleus; vf: ventral funiculus.

### Spreading of phosphorylated α-Syn from the lower urinary tract to the brain in TgM83^+/−^ mice via micturition reflex pathways

Our above-mentioned retrograde tracing study with FG has identified the pathways controlling EUS or DET. To demonstrate whether spreading of misfolded α-Syn via the same pathways induces the MSA-like neuropathology, we next injected α-Syn preformed fibrils (PFFs) (Figure 3-figure supplement 1) to EUS or DET (Figure 3-figure supplement 1) in TgM83^+/−^ mice and evaluated phosphorylated α-Syn in different sections at different time points.

**Figure 3.**
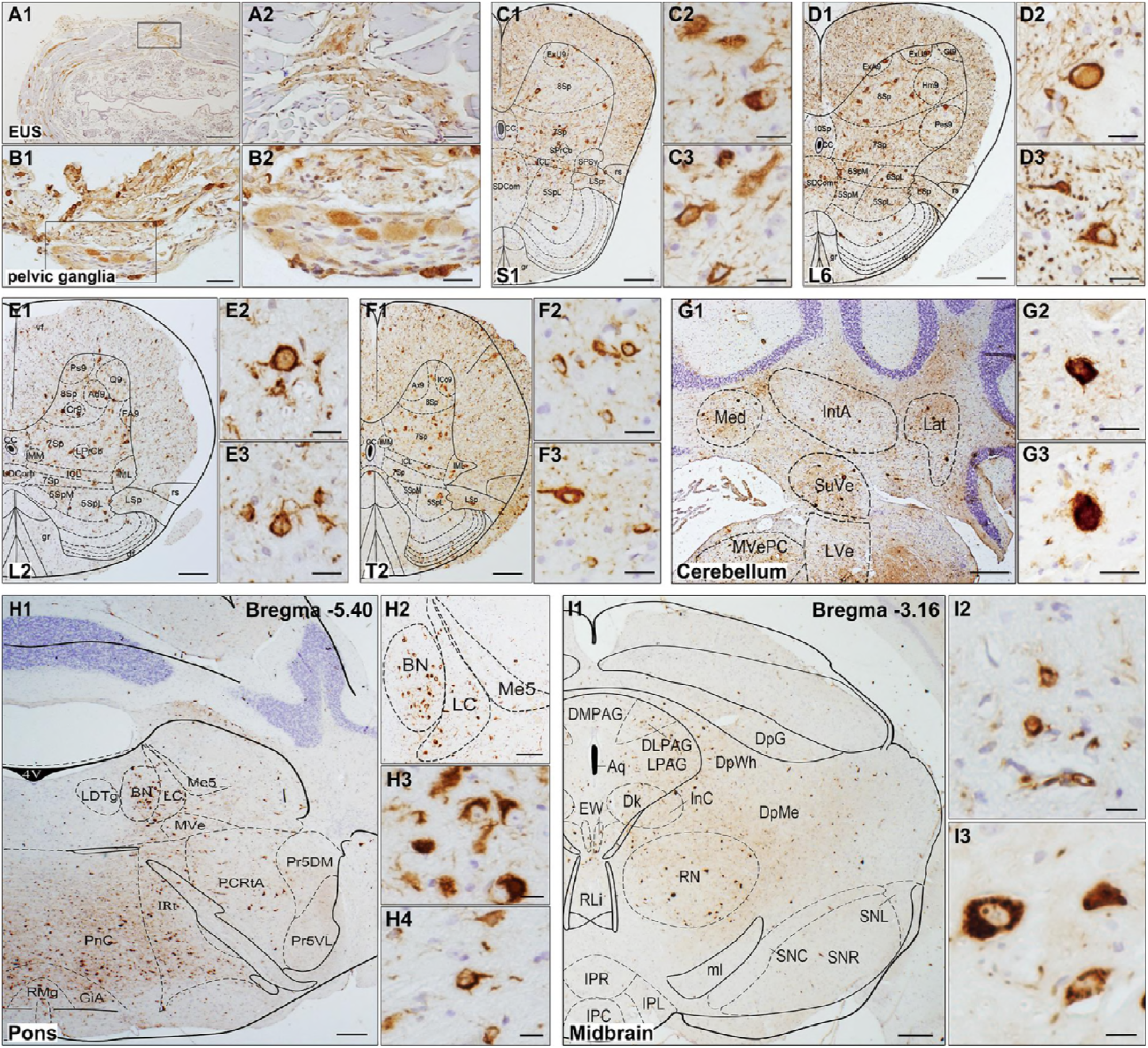
Representative immunohistochemical results of different segments from diseased EUS-α-Syn PFFs TgM83^+/−^ mice. (A-I) Pathological α-Syn stained with anti-phospho-α-Syn (Ser 129) antibody. Representative images displayed the distribution of pα-Syn in EUS (A1), pelvic ganglia (B1), S1 (C1), L6 (D1), L2 (E1), T2 (F1), cerebellum (G1), pons (H1), midbrain (I1). (A2-I2, C3-I3, H4) High-magnification views relative to the main image. [Scale bars, 500 µm (A1, G1, H2); 100 µm (A2, B1); 50 µm (B2, G2, I2, G3-I3, H4); 250 µm (C1-F1); 25 µm (C2-F2, C3-F3); 1 mm (H1, I1).] Abbreviation: DLPAG: dorsolateral periaqueductal gray; DMPAG: dorsomedial periaqueductal gray; DpG: deep gray layer of the superior colliculus; DpMe: deep mesencephalic nucleus; DpWh: deep white layer of the superior colliculus; EW: Edinger-Westphal nucleus; GiA: gigantocellular reticular nucleus; IntA: interposed cerebellar nucleus, anterior part; IPC: interpeduncular nucleus, caudal subnucleus; IPL: internal plexiform layer of the olfactory bulb; IPR: interpeduncular nucleus, rostral subnucleus; IRt: intermediate reticular nucleus; Lat: lateral (dentate) cerebellar nucleus; LDTg: tegmental nucleus; LPAG: lateral periaqueductal gray; LVe: lateral vestibular nucleus; Med: medial (fastigial) cerebellar nucleus; MVePC: medial vestibular nucleus; parvocellular part; MVe: medial vestibular nucleus; PnC: pontine reticular nucleus, caudal part; Pr5DM: principal sensory trigeminal nucleus, dorsomedial part; Pr5VL: principal sensory trigeminal nucleus, ventrolateral part; RLi: rostral linear nucleus of the raphe; RMg: raphe magnus nucleus; SuVe: superior vestibular nucleus.

Immunohistochemical results show that pα-Syn, immunostained with the anti-pα-Syn antibody, were detected at 5-month post-injection in both EUS-and DET-α-Syn PFFs TgM83^+/−^ mice (Fig. 3). In contrast, pα-Syn has not been detected in EUS-PBS, DET-PBS TgM83^+/−^, and C57BL/6 mice post-injection (Figure 3-figure supplement 2). In EUS-α-Syn PFFs TgM83^+/−^ mice, small numbers of pα-Syn were detected in EUS and pelvic ganglia (Fig. 3A, B). Furthermore, pα-Syn was detected at the S1, L6, L2, and T2 levels of spinal cord; and they mostly existed in laminae V-VII and IX of these levels (Fig. 3C-F). In brain, pα-Syn existed in pons and midbrain (Fig. 3H, I). These observations are consistent with the results of the above-mentioned FG study. Additionally, using immunohistochemical approach, pα-Syn was also found in the cerebellar nuclei (Fig. 3G). The neuropathological findings in DET-α-Syn PFFs TgM83^+/−^ mice were similar to those in EUS-α-Syn PFFs TgM83^+/−^ mice. In conclusion, transmission of pathological α-Syn in these mice invades not only the autonomic nervous system associated with urinary function, but also the extrapyramidal system via the micturition reflex pathways, and these findings are consistent with MSA pathology found in patient autopsy (Cykowski et al., 2015; Stemberger et al., 2010; VanderHorst et al., 2015; Yoshida, 2007). Thus, these findings suggest that pathological α-Syn spreads from the autonomic innervation of the lower urinary tract to extrapyramidal system via the micturition reflex pathways, leading to widespread α-Syn inclusion pathology.

To further characterize the nature of α-Syn-positive deposits in diseased EUS-or DET-α-Syn PFFs TgM83^+/−^ mice and the distribution of pathological α-Syn in various cells, double immunofluorescence staining was then employed. First, we revealed that deposits of pα-Syn colocalized with ubiquitin in spinal cord, cerebellum, and BN of diseased EUS-or DET-α-Syn PFFs TgM83^+/−^ mice (Fig. 4A, C, G). Then, we found that pα-Syn colocalized with Iba-1 (microglia marker) in spinal cord, ventral pons, and midbrain (Fig. 4B, J, K), which indicates that microglial activation is involved in the pathological process. Additionally, we observed a small amount of pα-Syn in LC (Fig. 4F) and a large number of pα-Syn in PAG and RN (Fig. 4H). No pα-Syn was detected in substantia nigra pars compacta (SNc) (Fig. 4H). We also noticed high level of ubiquitin protein in dopamine (DA) neurons of SNc (Fig. 4I). Finally, we identified that transmission of α-Syn PFFs through micturition reflex pathways resulted in axonal pathology and demyelination in diseased EUS-α-Syn PFFs TgM83^+/−^ mice with a distinct loss of myelin basic protein (MBP) and neurofilament (Fig. 4D, E). Taken together, these results suggest that injection of α-Syn PFFs into EUS or DET of TgM83^+/−^ mice initiates pathological α-Syn transmission, including microglia activation, axonal pathology, and demyelination.

**Figure 4.**
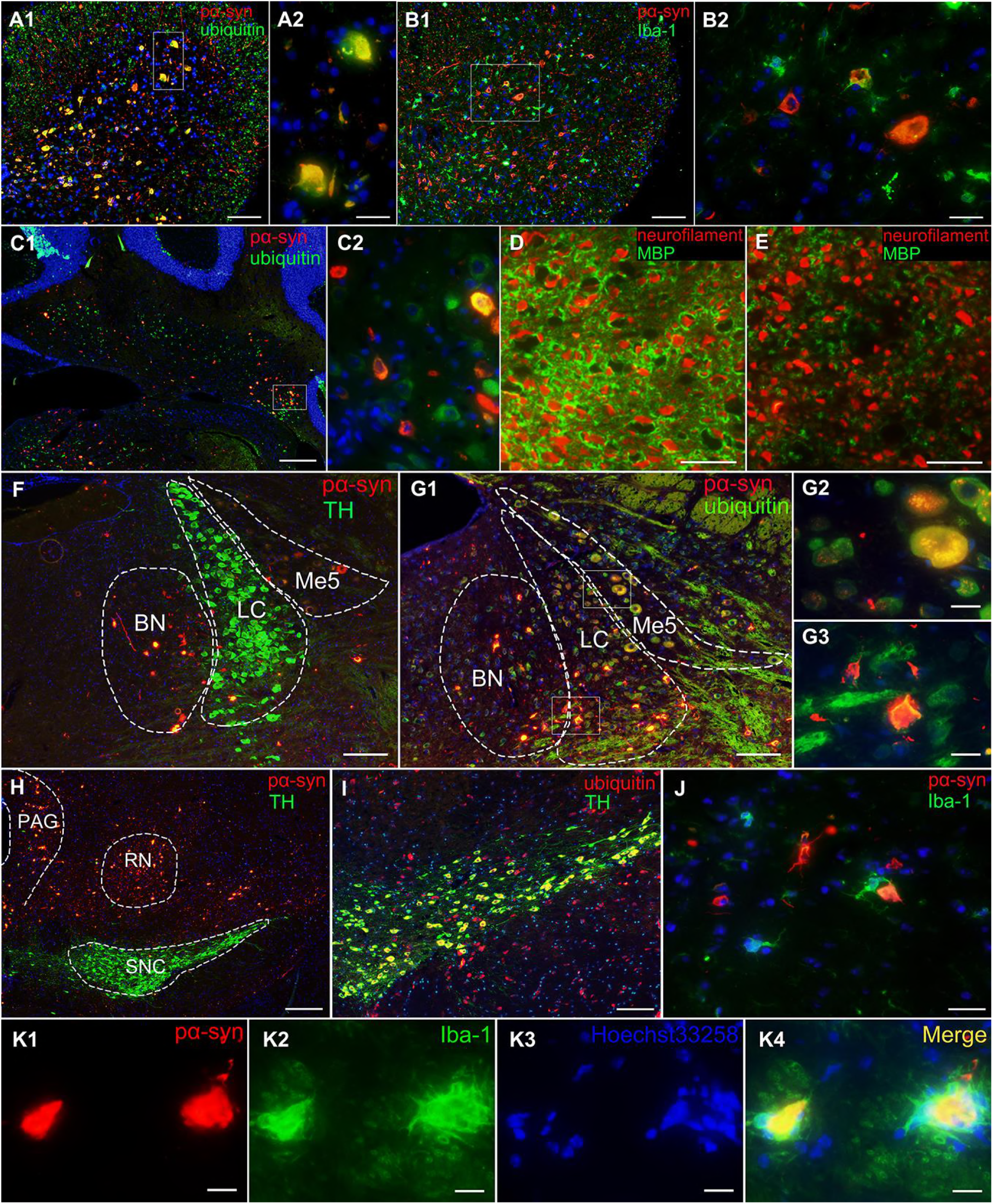
Double immunofluorescence analysis of different segments from diseased EUS-α-Syn PFFs TgM83^+/−^ mice and age-matched EUS-PBS TgM83^+/−^ mice. (A, B) Double immunofluorescence analysis of L6 from EUS-α-Syn PFFs TgM83^+/−^ mice for pα-Syn (red) and ubiquitin (green, A1-A2), Iba-1 (green, B1-B2). (C) Double immunofluorescence analysis of cerebellum from EUS-α-Syn PFFs TgM83^+/−^ mice for pα-Syn (red) and ubiquitin (green). (D, E) Double immunolabeling for MBP (green) and neurofilament (red) in L2 of EUS-PBS TgM83^+/−^ mice (D) and EUS-α-Syn PFFs TgM83^+/−^ mice (E). (F, G) Double immunofluorescence analysis of pons from diseased EUS-α-Syn PFFs TgM83^+/−^ mice for pα-Syn (red) and TH (green, F), ubiquitin (green, G1-G3). (H-J) Double immunofluorescence analysis of pα-Syn (red, H) and TH (green, H), ubiquitin (red, I) and TH (green, I), pα-Syn (red, J) and Iba-1 (green, J) in midbrain of diseased EUS-α-Syn PFFs TgM83^+/−^ mice. (K) Double immunolabeling for pα-Syn (red, K1) and Iba-1 (green, K2) in pons of EUS-α-Syn PFFs TgM83^+/−^ mice. Co-immunolabeling is represented by signal in yellow. Cell nuclei were counter stained with Hoechst33258 (blue). [Scale bars, 500 μm (A1, C1, H); 200 μm (B1, F, G1, I); 100 μm (G2-G3, K1-K4); 50 μm (A2, B2, C2, D, E, J).]

An immunoblot of spinal cord, pons, and PAG homogenate probed with α-Syn (Ser129P) and aggregated α-Syn (clone 5G4) antibodies was conducted to confirm that injection of α-Syn PFFs into EUS or DET of TgM83^+/−^ mice initiates MSA-like neuropathology (Fig. 5). In the insoluble fractions, immunostaining with α-Syn (Ser129P) reveals that bands around 15 kDa were detected in all examined diseased EUS-α-Syn TgM83^+/−^ mice, while it was faintly detected in EUS-α-PBS TgM83^+/−^ mice (Fig. 5A, D, F). Statistical analysis shows that pα-Syn is significantly increased in diseased EUS-α-Syn TgM83^+/−^ mice (Fig. 5B, E, G). Immunoblotting of the insoluble fractions of spinal cord homogenates using the aggregated α-Syn (clone 5G4) antibodies exhibits the bands around 35 kDa in diseased EUS-α-Syn TgM83^+/−^ mice but not in age-matched EUS-α-PBS TgM83^+/−^ mice (Fig. 5A). Statistical data indicate that the level of α-Syn aggregates is significantly elevated in EUS-α-Syn TgM83^+/−^ mice (Fig. 5C). These data indicate that the injection of α-Syn PFFs into EUS or DET can induce the spreading of pathological α-Syn in CNS.

**Figure 5.**
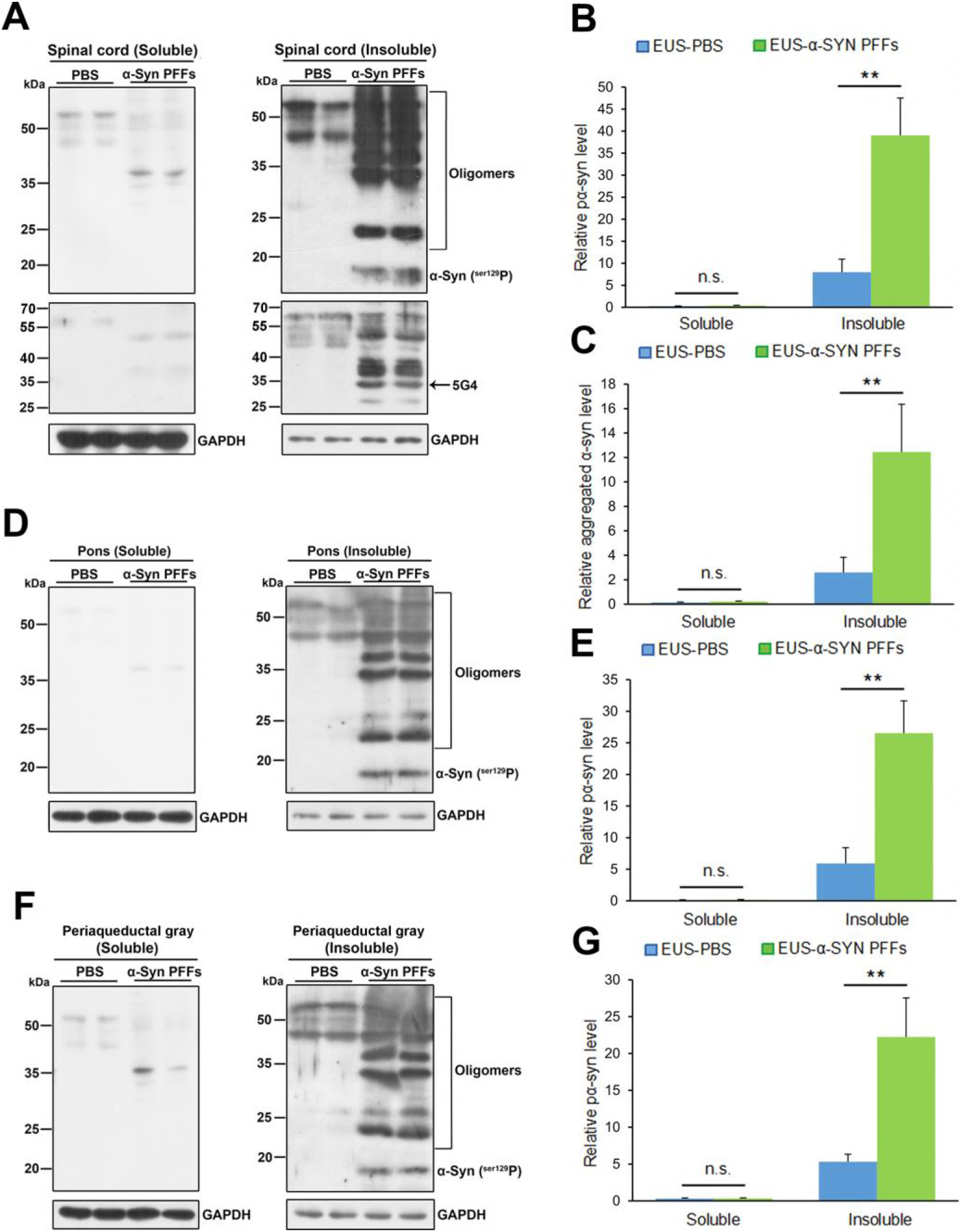
Western blot analysis of diseased EUS-α-Syn PFFs TgM83^+/−^ mice and age-matched EUS-PBS TgM83^+/−^ mice. (A, D, F) Representative immunoblots of α-Syn in the soluble and insoluble fractions of spinal cord, pons and PAG using the α-Syn (Ser129P) and aggregated α-Syn (clone 5G4) antibodies. Blots were probed for GAPDH as a loading control (Bottom). Molecular weight markers of migrated protein standards are expressed in kDa. (B, C, E, G) Quantification of soluble and insoluble α-Syn levels in the spinal cord, pons and PAG (n = 3 per group). Data are the means ± SD. Statistical significance was analyzed by using the Student’s t test and Mann-Whitney test, *P < 0.05; n.s., non-significant.

### Early-onset denervation-reinnervation of EAS in EUS-or DET-α-Syn PFFs TgM83^+/−^ and C57BL/6 mice

Forty age-matched healthy TgM83^*+/−*^ and C57BL/6 male mice were used; no abnormal EAS electromyogram (EMG) in these mice was detected. Based on the EMGs of the previous literature (Daube & Rubin, 2009; Palace, Chandiramani, & Fowler, 1997; Schwarz, Kornhuber, Bischoff, & Straube, 1997), abnormal EAS EMG would be defined if the EMG findings satisfied any one of the following six conditions: (1) fibrillation potentials; (2) positive sharp waves; (3) CRD; (4) fasciculation potentials; (5) myokymic discharges; and (6) satellite potential.

EUS-and DET-α-Syn PFFs TgM83^+/−^ mice show abnormal EAS EMGs at 2-month post-injection (P < 0.05), while no abnormality of EAS EMG was detected in PBS groups. Representative abnormal and normal EAS EMGs are shown in Fig. 6A-E and F, respectively. The prevalence of abnormal EAS EMGs in EUS-α-Syn PFFs TgM83^+/−^ mice was 55%, 84%, and 90% at 2-month, 4-month, and 6-month post-injection, respectively, versus 51%, 83%, and 91% in DET-α-Syn PFFs TgM83^+/−^ mice, respectively (Fig. 6G, H). In C57BL/6 mice, the data of abnormal EAS EMGs show no significant difference between EUS-or DET-α-Syn PFFs groups and PBS groups (Fig. 6I, J). The results suggest prevalence of abnormal EAS EMGs increases along with the progression of neural lesions caused by α-Syn PFFs. We also injected α-Syn PFFs into the intestine wall of stomach and duodenum of TgM83^+/−^ mice. However, the TgM83^+/−^ mice with intestine-α-Syn PFFs didn’t develop abnormal EAS EMG while TgM83^+/−^ mice did. Taken together, the denervation-reinnervation of EAS occurs in the early stage of neuropathological process in a time dependent manner, and may be caused by spreading of α-Syn PFFs from the lower urinary tract through micturition reflex pathways.

**Figure 6.**
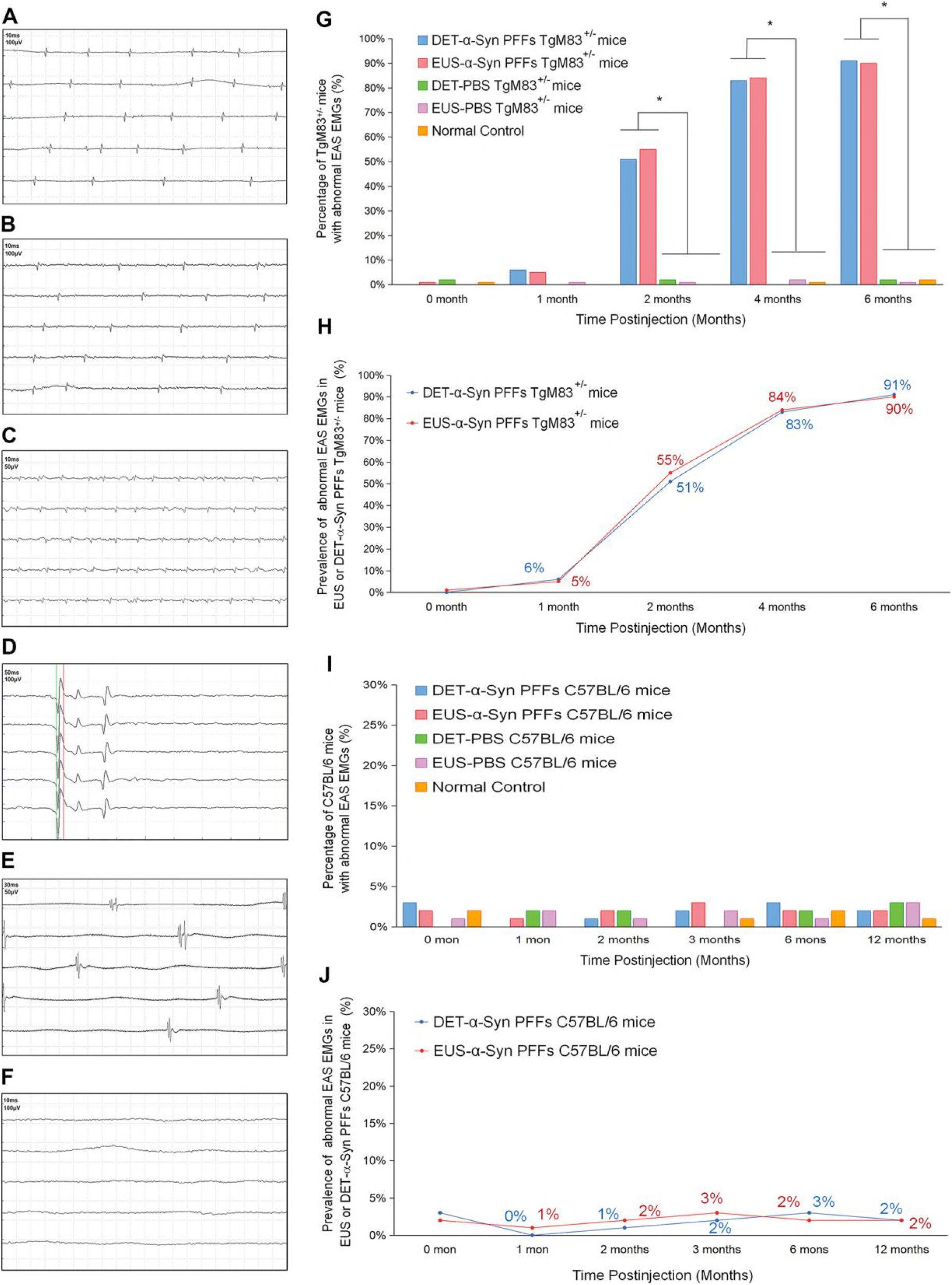
EAS EMG analysis of TgM83^+/−^ mice and C57BL/6 mice. (A-E) Representative abnormal EAS EMGs from diseased EUS-or DET-α-Syn PFFs TgM83^+/−^ mice. Abnormal EAS EMGs show as fibrillation potential (A), positive sharp waves (B), CRD (C), satellite potential (D), and myokymic discharges (E). (F) Representative normal EAS EMG referring to resting potential from EUS-PBS TgM83^+/−^ mice at 5-month post-injection. (G-J) Time-dependent distribution of abnormal EAS EMGs in different groups of TgM83^+/−^ mice (G, H) and C57BL/6 mice (I, J). EUS-α-Syn PFFs mice n = 20, DET-α-Syn PFFs mice n = 18, EUS-PBS mice n = 18, DET-PBS mice n = 16, control group n = 20. Statistics was analyzed employing the χ2 test. *P < 0.05 relative to the corresponding PBS groups and control groups.

### Urinary dysfunction in EUS-or DET-α-Syn PFFs TgM83^+/−^ mice

The urodynamic baseline is determined by cystometry results of 2-month-old male TgM83^+/−^ and C57BL/6 mice prior to treatments. Urinary dysfunction was observed in EUS-and DET-α-Syn PFFs TgM83^+/−^ mice between 3 and 4 months post-injection and persisted to the last stage examined. At 4-month post-injection, both EUS-and DET-α-Syn PFFs TgM83^+/−^ mice exhibited a significant increase in amplitude, PVR, and NVCs during the filling phase compared to PBS groups (P < 0.05). Meanwhile, VV and ICI in EUS-or DET-α-Syn PFFs TgM83^+/−^ mice were found less and shorter, respectively (Fig. 7). The body mass of EUS-or DET-α-Syn PFFs TgM83^+/−^ mice was mostly lighter than EUS-or DET-PBS TgM83^+/−^ mice; however, the bladder of EUS-or DET-α-Syn PFFs TgM83^+/−^ mice exhibited overtly greater size compared to EUS-or DET-PBS mice (Figure 7-figure supplement 1), which was probably due to progressive urothelium and DET hyperplasia in α-Syn PFFs TgM83^+/−^ mice. By the time of the 14th month post-injection, EUS-or DET-α-Syn PFFs C57BL/6 mice did not show any urinary dysfunction. The intestine-α-Syn PFFs TgM83^+/−^ mice didn’t show any urinary dysfunction at 3.5-month post-injection neither when EUS-and DET-α-Syn PFFs TgM83^+/−^ mice did already. All results mentioned above suggest that urodynamic assessment in EUS-or DET-α-Syn PFFs TgM83^+/−^ mice was characterized by an overactive, less stable, and inefficient bladder. In addition, α-Syn PFFs injection into EUS or DET of TgM83^+/−^ mice caused potential dyssynergia between DET and EUS, leading to hyperactive bladder and DET hyperreflexia (Boudes et al., 2013; Hamill et al., 2012), which resembles urinary dysfunction in patients with MSA. Thus, we developed an animal model to replicate MSA-like urinary disorders and abnormal EAS EMGs, which has not been previously reported.

**Figure 7.**
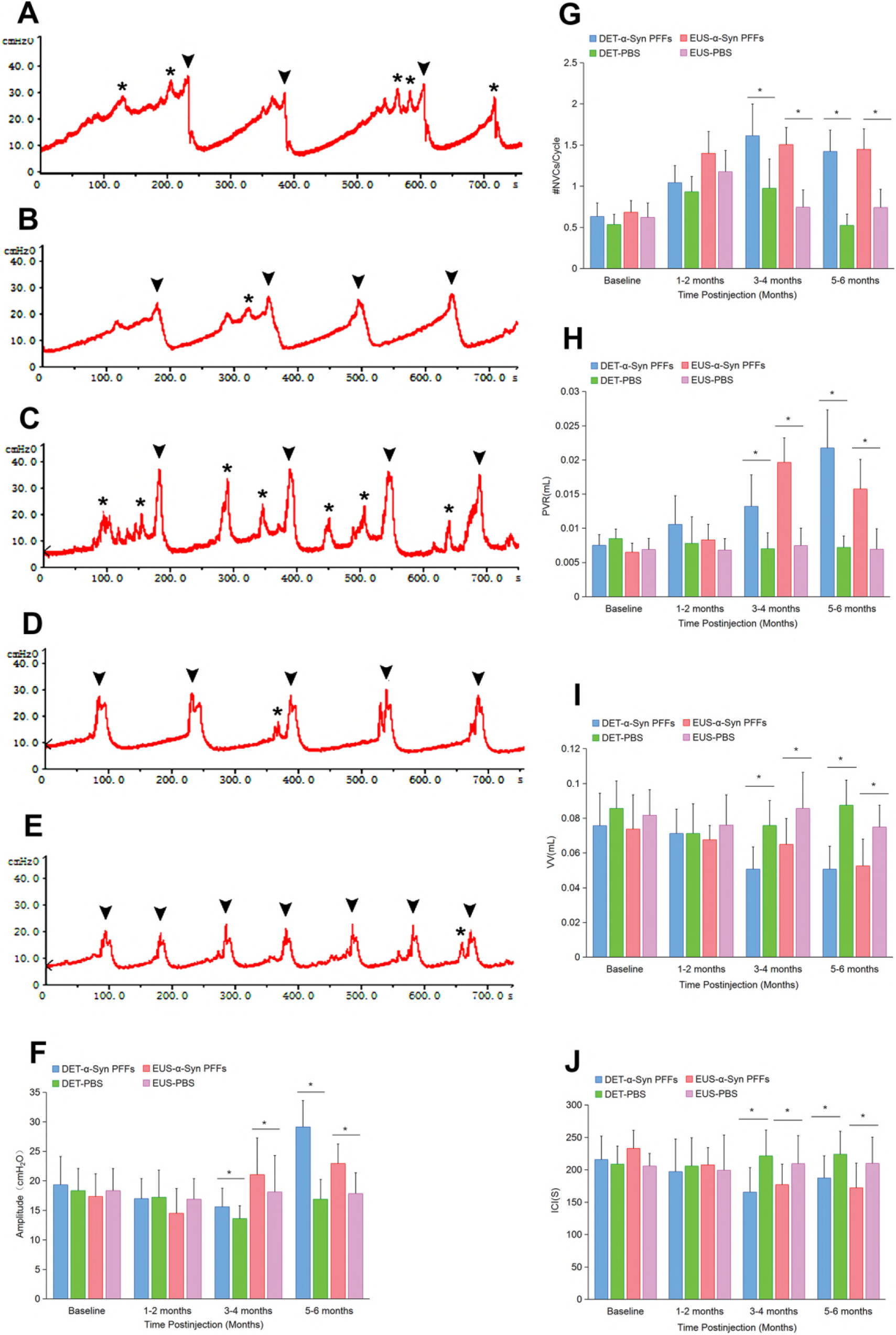
Urinary function analysis of TgM83^+/−^ mice. (A-E) Representative cystometry traces in DET-α-Syn PFFs (A), DET-PBS (B), EUS-α-Syn PFFs (C), EUS-PBS (D) TgM83^+/−^ mice at 5-month post-injection and baseline group (E). Arrows indicate void events and asterisks indicate NVCs. (F-J) Summary bar graphs from urodynamic evaluation for EUS and DET TgM83^+/−^ mice including amplitude (F), #NVCs/Cycle (G), PVR (H), VV (I) and ICI (J). EUS-α-Syn PFFs TgM83^+/−^ mice n = 18, DET-α-Syn PFFs TgM83^+/−^ mice n = 16, EUS-PBS TgM83^+/−^ mice n = 22, DET-PBS TgM83^+/−^ mice n = 20. Data are the means ± SD. Statistics was analyzed employing the Student’s t test and Mann-Whitney test. *P < 0.05 indicates a significant difference between EUS-or DET-α-Syn PFFs groups and EUS-or DET-PBS groups.

### Motor impairments in EUS-or DET-α-Syn PFFs TgM83^+/−^ mice

Both EUS-and DET-α-Syn PFFs TgM83^+/−^ mice began to exhibit motor impairments from 5-month post-injection. Most diseased mice presented an arched back initially and then progressed with weight loss, ataxia, paralysis, and a moribund state requiring euthanasia within 3 weeks (Fig. 8A). Compared with DET-α-Syn PFFs TgM83^+/−^ mice, the behavioral deficiency was more obvious in EUS-α-Syn PFFs TgM83^+/−^ mice. At 5-month post-injection, α-Syn PFFs TgM83^+/−^ mice showed significantly increased motor behavioral scale (MBS) score compared with EUS-or DET-PBS TgM83^+/−^ mice, which is considered as a semi-quantitative assessment for MBS rating (Fig. 8B and Figure 8-figure supplement 1). The rotarod test was carried out to assess coordination capability. The performance on the rotating rod was significantly impaired in EUS-and DET-α-Syn PFFs TgM83^+/−^ mice compared to PBS controls, as their latency to fall was markedly reduced (Fig. 8C). In an open field test, EUS-and DET-α-Syn PFFs TgM83^+/−^ mice showed significantly reduced spontaneous activities in comparison with PBS-injected mice (Fig. 8J, K). Footprint analysis indicates that EUS-and DET-α-Syn PFFs TgM83^+/−^ mice have shorter stride length and wider base width compared to PBS-injected mice (Fig. 8D, E and Figure 8-figure supplement 1). Moreover, EUS-and DET-α-Syn PFFs TgM83^+/−^ mice also showed significantly motor dysfunction in the beam walking test (Fig. 8F, G) and pole test (Fig. 8H, I). EUS-or DET-PBS TgM83^+/−^ mice didn’t show any phenotype until they were 22 months old, consistent with our spontaneously sick TgM83^+/−^ mice in timeline. As the previous study reported, spontaneously sick TgM83^+/−^ mice develop series of phenotypes between 22-28 months of age (Giasson et al., 2002). Nevertheless, EUS-and DET-α-Syn PFFs C57BL/6 mice failed to exhibit behavioral abnormalities up to 420-day post-injection (Figure 8-figure supplement 2).

**Figure 8.**
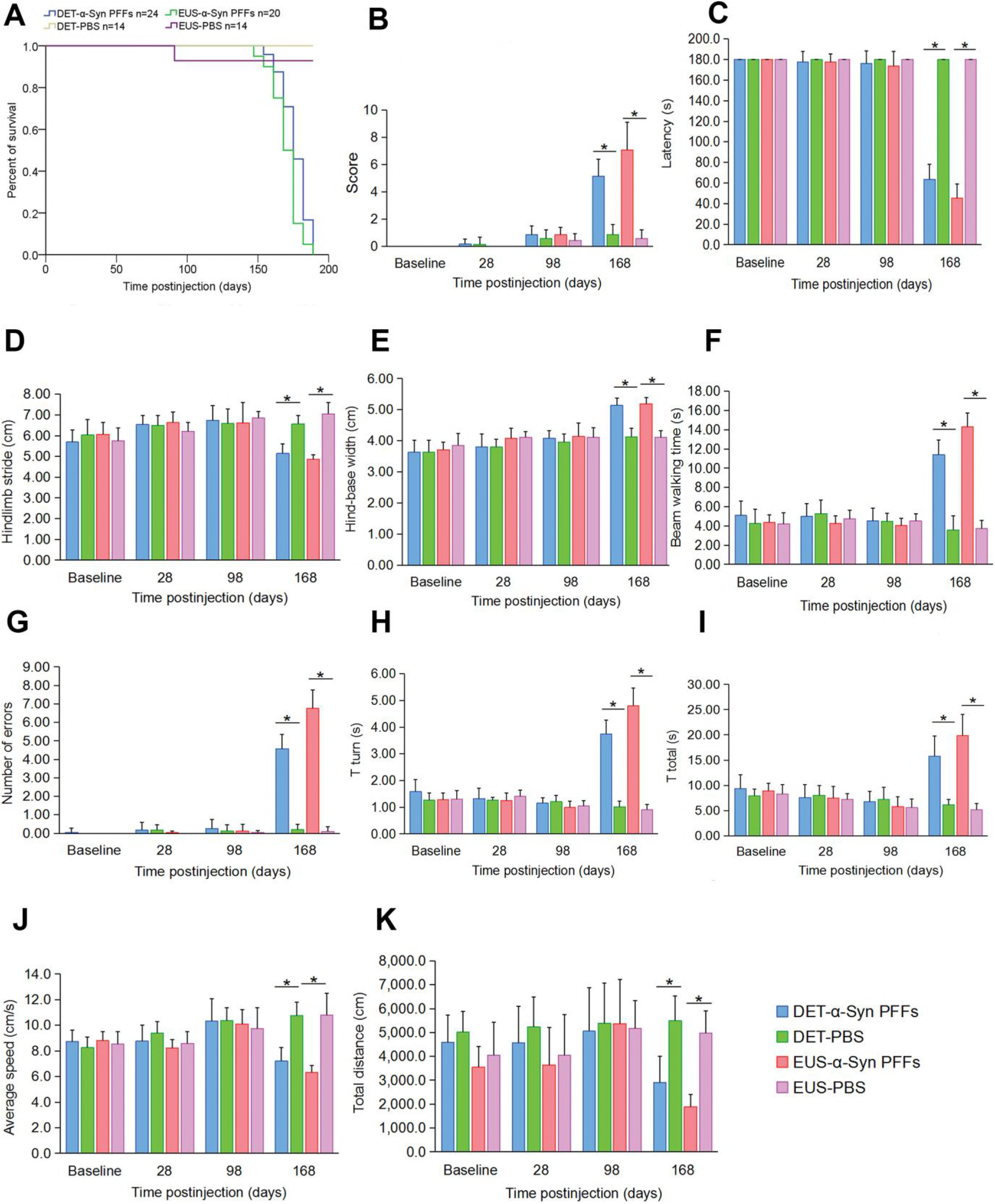
Behavioral analysis of TgM83^+/−^ mice. (A) Kaplan-Meier survival plot shows decreased survival time (due to death or euthanasia because of paralysis) for α-Syn PFFs TgM83^+/−^ mice compared with age-matched PBS TgM83^+/−^ mice. (B) The mean score of MBS. (C) Latency to fall from the rotarod. (D and E) Footprint analysis of the hindlimb stride length (D) and the hind-base width (E). (F and G) The average time to cross the beam (F) and the average number of side slip errors (G) on the beam. (H and I) T turn (H) and T total (I) of the pole test. (J and K) Average speed (J) and total distance (K) traveled during 15 minutes in the open field test. EUS-α-Syn PFFs TgM83^+/−^ mice n = 12, DET-α-Syn PFFs TgM83^+/−^ mice n = 20, EUS-PBS TgM83^+/−^ mice n = 14, DET-PBS TgM83^+/−^ mice n = 14. Data are the means ± SD. Statistical analysis was done by using the Student’s t test and Mann-Whitney test, *P < 0.05.

Taken together, EUS-and DET-α-Syn PFFs TgM83^+/−^ mice developed distinct motor signs including weight loss, bradykinesia, ataxia, and paralysis at 5-month post-injection. We conclude that injection with α-Syn PFFs into EUS or DET in TgM83^+/−^ mice initiates MSA-like motor deficits.

## Discussion

We broadened the landscape of the pathogenesis of synucleinopathies. Prior to this study, we had some understanding of the pathogenesis of PD, while we knew less about that of MSA. In MSA, autonomic dysfunctions, especially urinary dysfunction (Kirby, Fowler, Gosling, & Bannister, 1986), are severe and common which are different from the other main synucleinopathy, e.g., PD. Based our long-term observations of clinical subjects with these diseases, we reckoned that pathological α-Syn might exist in the lower urinary tract at the early stage of MSA instead of gut as shown in PD (Holmqvist et al., 2014). To test this hypothesis, we first performed bladder biopsy in participants. The findings from this study show that misfolded α-Syn aggregates indeed exist in DET or EUS in 71.9% of the included MSA patients. The subsequent results from immunohistochemistry studies in experimental mice show that α-Syn aggregates invade the micturition reflex pathways. In addition, we found widely positive staining of pα-Syn in ventral white matter of spinal cord, possibly due to nerve tracts from brain and comprehensively longitudinal connections by synapses of numerous nerve fibers. Moreover, we detected overt α-Syn aggregates in cerebellar nucleus which indicate that α-Syn aggregates transmit to cerebellum via rubro-cerebello-rubrospinal circuit (Larson-Prior & Cruce, 1992). α-Syn pathology in EUS-or DET-α-Syn PFFs C57BL/6 mice was not detected at 14-month post-injection. The results of double immunofluorescence analysis further demonstrate the pathological lesions in CNS of the two mouse models. There was apparent microglial activation and demyelination in CNS of EUS-α-Syn PFFs TgM83^+/−^ mice, which is a major pathological feature of MSA (Ettle et al., 2016). The immunostaining results validate the hypothesis that pathological α-Syn transmits initially from urogenital autonomic nerves to extrapyramidal system, inducing α-Syn inclusion pathology. EAS EMG has been previously proposed as a diagnostic method for MSA (E. A. Lee, Kim, & Lee, 2002). Abnormalities of EAS EMG in MSA indicate the denervation-reinnervation of EAS caused by neuronal loss of Onuf’s nucleus in the anterior horn of the spinal cord (E. A. Lee et al., 2002; Libelius & Johansson, 2000). In this study, we conducted EAS EMG in mouse models to assess denervation-reinnervation of EAS. Remarkably, abnormal EAS EMGs emerged at 2-month post-injection in EUS-or DET-α-Syn PFFs TgM83^+/−^ mice. Further, overall prevalence reached 90% in EUS-α-Syn PFFs TgM83^+/−^ mice versus 91% in DET-α-Syn PFFs TgM83^+/−^ mice at 6-month post-injection, while no abnormality was detected in PBS-injected TgM83^+/−^ mice. We demonstrate that electromyography experimental results in the mouse models are similar to EAS EMG feature of MSA patients preceding urinary dysfunction and movement disorders. Previous studies (Yamamoto et al., 2005) presented a view that selective neuronal loss of Onuf’s nucleus, which innervates EAS, results in abnormal EAS EMGs in patients with MSA. We found that α-Syn aggregates are present in Onuf’s nucleus in both EUS-and DET-α-Syn PFFs TgM83^+/−^ mice.

In this study, we implemented urodynamic assessment in different time points to evaluate urinary function in experimental mice. Consequently, EUS-or DET-α-Syn PFFs TgM83^+/−^ mice exhibited urodynamic changes after the 3rd month post-injection, prior to motor impairments, versus no changes in PBS groups. Here, we identified that urinary dysfunction, characterized by urinary bladder hyperreflexia of α-Syn PFFs TgM83^+/−^ mice, replicates the altered bladder function in MSA patients including urinary incontinence, frequency, urgency, and retention (Fowler, Dalton, & Panicker, 2010; Ragab & Mohammed, 2011). Previous study (Libelius & Johansson, 2000) showed that the spontaneous TgM83^+/−^ mice developed urinary bladder dysfunction prior to motor dysfunction due to A53T mutant α-Syn. In our study, EUS-or DET-α-Syn PFFs TgM83^+/−^ mice started to perform urinary dysfunction at 3.5-month post-injection whereas EUS-or DET-PBS and non-inoculated TgM83^+/−^ mice didn’t show any urinary dysfunction until they were 22 months old. The occurrence of urinary dysfunction in EUS-or DET-α-Syn PFFs TgM83^+/−^ mice is earlier than PBS control groups and non-inoculated TgM83^+/−^ mice. In our α-Syn PFFs TgM83^+/−^ mice, the micturition reflex pathways, including EUS and DET, pelvic ganglia, Onuf’s nucleus, IML, PAG, BN, and LC, exhibited misfolded α-Syn aggregates, revealing pathological mechanisms of the urinary dysfunction. However, we did not observe appreciable levels of misfolded α-Syn deposition in oligodendrocytes within the TgM83^+/−^ mice. This observation could be explained by misfolded α-Syn originating from different parts of PNS to CNS via neuronal projections transsynaptically. In spontaneously ill TgM83^+/−^ mice, α-Syn aggregates have not been detected in ExU9, ICL, ExA9, Gl9, sacral dorsal commissural nucleus, IML, LDCom, BN, and PAG, which is different from the diseased EUS-α-Syn PFFs TgM83^+/−^ mice. As these spared areas are involved in controlling the urinary bladder (Fowler et al., 2008), these data support that the preceding autonomic dysfunction of EUS-or DET-α-Syn PFFs TgM83^+/−^ mice results from exogenously injected α-Syn PFFs instead of A53T mutant α-Syn. Therefore, our results suggest that misfolded α-Syn spreading through the micturition reflex pathways retrogradely may lead to urinary dysfunction.

Previous studies indicate that pathological α-Syn spread from peripheral nervous system (PNS) to CNS through retrograde axonal transport, in a stereotypical and topographical pattern (Bernis et al., 2015; Braak et al., 2003; Holmqvist et al., 2014; Luk, Kehm, Carroll, et al., 2012). Experimental studies suggest that PD pathology may originate in the vagal nerves from the gut and gradually propagate to the brain (Holmqvist et al., 2014). These findings support the hypothesis that different synucleinopathies may originate from different part of PNS and gradually propagate to CNS. Hence, we speculate that pathological α-Syn originate from the autonomic innervation of the lower urinary tract has the potential to propagate to CNS and induce MSA Our data demonstrate that misfolded α-Syn can induce α-Syn inclusion pathology along with autonomic failure and motor impairments by transmitting from the autonomic control of the lower urinary tract to the brain via micturition reflex pathways.

As previously reported by others, peripheral injection of α-Syn PFFs into multiple sites could promote the development of α-Syn pathology in the CNS of TgM83^+/−^ mice (Ayers et al., 2017; Breid et al., 2016; Holmqvist et al., 2014; Luk, Kehm, Carroll, et al., 2012; Sacino, Brooks, Thomas, McKinney, Lee, et al., 2014; Sacino, Brooks, Thomas, McKinney, McGarvey, et al., 2014; Watts et al., 2013). We injected α-Syn PFFs into the striatum of TgM83^+/−^ mice in initial studies. The results show that the lower urinary tract pathology can’t be obtained at 6th month after intracerebral α-Syn PFFs injection. We also injected α-Syn PFFs into the intestine wall of stomach and duodenum of TgM83^+/−^ mice. However, the intestine-α-Syn PFFs TgM83^+/−^ mice didn’t develop abnormal EAS EMG and urinary dysfunction at 5-month post-injection when they had α-Syn pathology in CNS and motor impairments already. Thus, α-Syn injection into intestine wall alone couldn’t induce the denervation-reinnervation of EAS and urinary dysfunction prior to motor impairments in TgM83^+/−^ mice. Again, this further indicates that α-Syn of PD and MSA may start in different places.

Here we further demonstrate that injection with α-Syn PFFs into EUS or DET induces a rapid progression of motor dysfunctions. Our study shows that injection with α-Syn PFFs into EUS or DET in TgM83^+/−^ mice causes not only seeding of α-Syn aggregation in the CNS, but also rapid progressive motor dysfunctions evaluated using a spectrum of behavioral tests. From our findings, the occurrence of motor impairments in our EUS-or DET-α-Syn PFFs TgM83^+/−^ mice was much earlier than spontaneously ill TgM83^+/−^ mice (Giasson et al., 2002). According to previous studies (Breid et al., 2016; Holmqvist et al., 2014; S. B. Prusiner et al., 2015; Sacino, Brooks, Thomas, McKinney, Lee, et al., 2014; Sacino, Brooks, Thomas, McKinney, McGarvey, et al., 2014), the animal models of synucleinopathy induced by exogenous inoculation involve different inocula and inoculation positions, developing variable α-Syn pathology and motor impairments without autonomic dysfunction. Furthermore, pathology of α-Syn inclusions observed in motor neuron of ventral horn, cerebellum, and RN in α-Syn PFFs TgM83^+/−^ mice provides compelling neuropathological evidence for the motor impairments.

In summary, this study suggests one possible pathogenic mechanism of MSA, which is the spreading of α-Syn inclusion pathology from the autonomic control of the lower urinary tract to the brain. Also, our data support the view that pathological α-Syn may originate from different parts of PNS among different disorders of synucleinopathies. However, the pathogenic mechanisms of MSA are not fully understood, other possibilities may exist. Thus, we and others will need to investigate further.

## Materials and methods

### Patients

Forty-five patients (18 men, 27 women; age 60.6 ± 7.2 years) were enrolled consecutively from 2016 to 2018 with MSA (32 patients), PD (7 patients), or progressive supranuclear palsy (PSP) (6 patients) according to consensus criteria (Kalia & Lang, 2015; Litvan et al., 1996; Stefanova et al., 2009), respectively. In MSA, the phenotype was characterized by prevalently cerebellar signs in 19 patients and by parkinsonian signs in the remaining 13 patients. Disease severity was evaluated using Unified Multiple System Atrophy Rating Scale (UMSARS) (Low et al., 2015; Wenning et al., 2004). UMSARS Total is a sum of UMSARS Ⅰ and UMSARS Ⅱ. Demographic and clinical data are summarized in Table 1. At the time of enrollment, all subjects underwent clinical and electrophysiological evaluation as well as EUS and bladder biopsies at 3 sites: left wall, right wall, and triangle region. Twenty subjects were also included in the study as controls (7 men, 13 women; age 58.5 ± 7.0 years). All biopsies were performed according to the outpatient procedures by experienced urologists in a prescriptive exam room. Cystoscopy was performed using standard cystoscope according to previously published procedures under local anesthesia with 1% xylocaine (Butros, McCarthy, Karaosmanoglu, Shenoy-Bhangle, & Arellano, 2015). The procedure was repeated until the EUS, left wall, right wall, and triangle region of bladder tissues were obtained. Samples were immediately fixed in 4% paraformaldehyde and kept at 4 ℃ for at least 2 days. The study was executed with the approval of the Institutional Ethics Committees of the Zhengzhou University.

### Animals

TgM83^+/−^ and C57BL/6 mice were purchased from Nanjing Biomedical Research Institute of Nanjing University (Nanjing, China), and evaluated at the age of six to eight weeks. The hemizygous TgM83 mice expressed the human A53T α-Syn driven by the prion gene promoter (Giasson et al., 2002). C57BL/6 mice were chosen as the control mice because TgM83^+/−^ mice were maintained on a mixed C57/C3H genetic background. Mice were kept in a near pathogen-free environment under standard conditions with food and water (21 °C, 12h/12h light-dark cycle). All experiments were conducted in accordance with the Guide for the Care and Use of Laboratory Animals. The protocols were approved by the Institutional Ethics Committees of the Zhengzhou University.

### α-Synuclein preformed fibrils (PFFs) preparation

α-Syn (S-1001, rPeptide) was resuspended in assembly buffer (20 mM Tris-HCl, 100 mM NaCl, pH 7.4) at concentration of 1 mg/ml. To obtain PFFs, the samples were placed in 2 ml sterile polypropylene tubes, sealed with parafilm, and agitated in a beaker with a magnetic stirrer (MS-H-Pro+, Scilogex, China) at 350 rpm for 7 days at 37 °C. After 7 days of incubation, the α-Syn fibrils were sonicated for 45 seconds using an ultrasonic cell disruptor at 10% of its peak amplitude (Scientz-IID, Ningbo, China). α-Syn fibrils were stored at −80°C until use.

### Modeling surgery

All injections were performed using a manual microinjector under an operating microscope. Mice were anesthetized with isoflurane inhalation and fixed in a supine position. After disinfection, mice were inoculated in the EUS with 15 μl α-Syn PFFs (1 mg/ml) or phosphate buffered saline (PBS, Solarbio), or DET with 20 μl (1 mg/ml) α-Syn PFFs or PBS. Following the injection, the exposed wound was sewn closed.

### EAS EMG

EAS EMG was carried out in all animals before injection for control groups and at correspondingly post-injection times to determine EAS denervation-reinnervation. Animals in each group underwent EAS EMG as follows (Aghaee-Afshar et al., 2009; Buffini, O’Halloran, O’Herlihy, O’Connell, & Jones, 2012; Healy, O’Herlihy, O’Brien, O’Connell, & Jones, 2008; Lane et al., 2013): anesthesia was induced using isoflurane inhalation. Limb withdrawal to paw pinch and corneal reflexes of animals were observed to assess the level of anesthesia. After placing the animal supine, shaving the thigh, and establishing a ground connection, a disposable concentric 30-gauge needle electrode (Technomed Europe), which has a 25-mm length, 0.30-mm diameter, and 0.021-mm^2^ recording area, was inserted at the 3 or 9 o’clock position of the anal orifice perpendicularly into the EAS from the perianal skin close to the mucocutaneous junction to a depth of approximately 1 to 2 mm. The point of the electrode insertion was adjusted under audio guidance until a permanent tonic activity was recorded, in order to ensure that the electrode has entered EAS. If the mouse discharged a fecal pellet during the recording process, a pair of forceps were used to gently clip it out. EMG was performed with an EMG monitoring machine (MEB-2306C, NIHON KOHDEN CORPORATION, Tokyo, Japan) at a sweep speed of 10 ms/div and a gain of 100 uv/div. Abnormal EAS EMGs were simultaneously visualized and recorded. EMG activity was quantified by prevalence of abnormal EAS EMGs (fibrillation potentials, positive sharp waves, complex repetitive discharges (CRD), fasciculation potentials, myokymic discharges, and satellite potential) in each group (Daube & Rubin, 2009; Palace et al., 1997; Schwarz et al., 1997).

### Cystometry evaluations and calculations

The following urodynamic parameters (Boudes et al., 2013; Fandel et al., 2016; Girard, Tompkins, Parsons, May, & Vizzard, 2012; Y. S. Lee et al., 2013; Silva et al., 2015) were used for the current study: (1) Maximum voiding pressure (P_max_; cmH_2_O); (2) Basal bladder pressure (BBP; cmH_2_O): the lowest bladder pressure during filling phase; (3) Amplitude (cmH_2_O): P_max_ - BBP; (4) Bladder leak point pressure (BLPP; cmH_2_O): intravesical pressure recorded at the first leaking/micturition point; (5) Threshold pressure (Pt; cmH_2_O); (6) Nonvoiding contractions during filling phase (NVCs): rhythmic intravesical pressure rises (> 5 cmH_2_O from baseline pressure) without any fluid leakage from the urethra; (7) Postvoid residual volume (PVR; ml): the remaining saline in the bladder collected and measured after stopping the infusion at the end of the final micturition cycle; (8) Maximum bladder capacity (MBC; ml): volume between the start of infusion and the BLPP; (9) Voided volume (VV; ml): MBC - PVR; (10) Intercontraction interval (ICI; s).

### Statistical analysis

All statistical analyses were performed using SPSS 21.0 (IBM, Armonk, New York, USA). Characteristics of patients presented as mean ± SD, statistical differences among groups of subjects were assessed using χ^2^ test. Additionally, behavioral data and cystometry parameters of mice were presented as mean ± SD, employing Student’s t test for comparison between two groups while one-way ANOVA for three when these data were distributed normally (P > 0.05 by Shapiro-Wilk test). Otherwise, the Mann-Whitney test was used for two groups versus Kruskal-Wallis test for three. Comparative analysis for prevalence of abnormal EAS EMGs among groups was performed by means of χ^2^ test. Pvalues < 0.05 were considered to be statistically significant.

## Acknowledgements

This work was supported by grants from the National Natural Science Foundation of China (No. 81671267, 81471307, 81873791, 81301086, and 81430023). We thank Dr. Richard M. Niles for his thoughtful editing of the manuscript.

## Supplementary Materials

**Figure 3-figure supplement 1.**
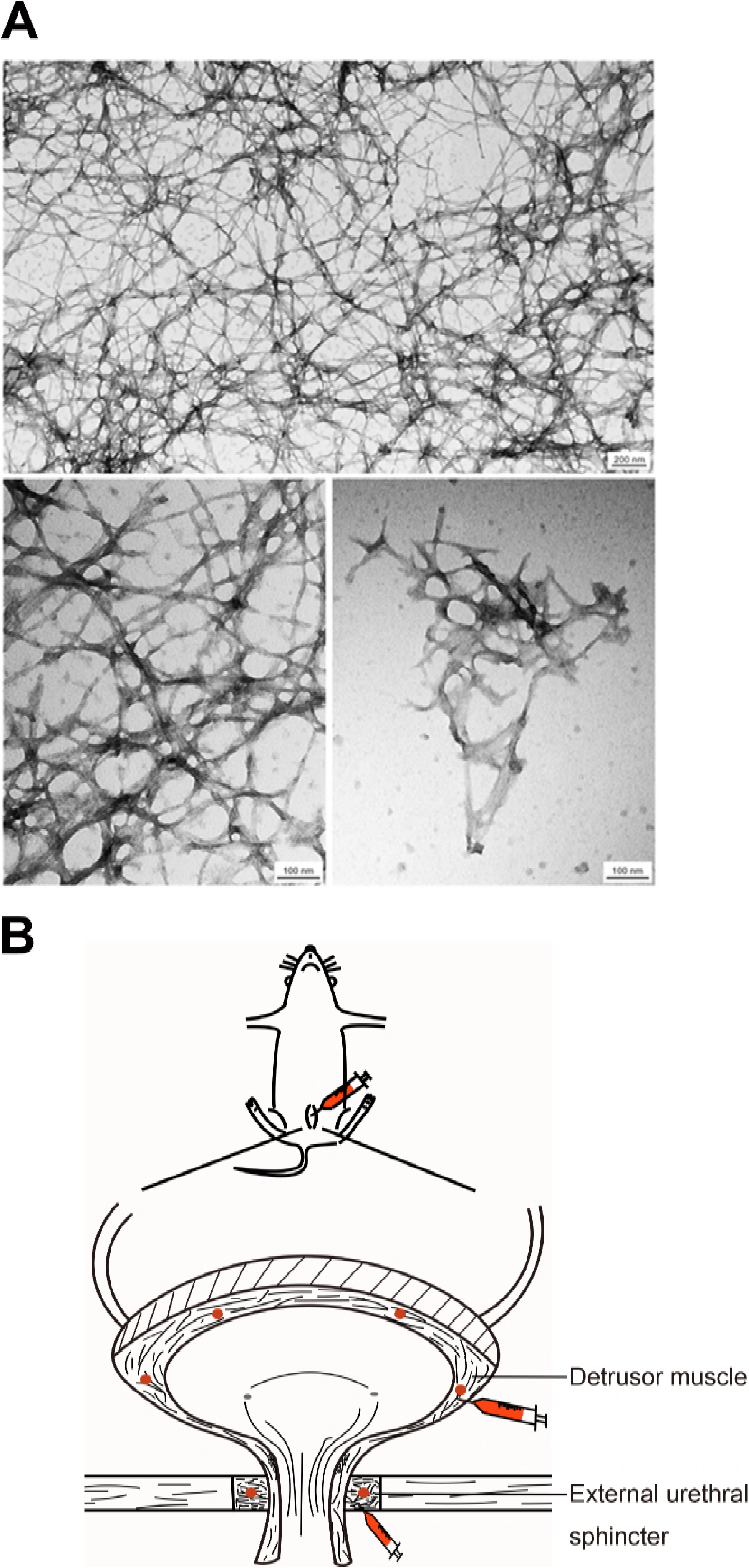
(**A**). Negatively stained transmission electron micrographs of α-Syn PFFs (upper), α-Syn PFFs in a high magnification (lower left), and α-Syn PFFs which were sonicated and used for modeling (lower right). (**B**) Schematic displayed the positions where were inoculated by α-Syn PFFs, including DET and EUS. [Scale bars, 200 nm (a, upper); 100 nm (a, lower left and right).]

**Figure 3-figure supplement 2.**
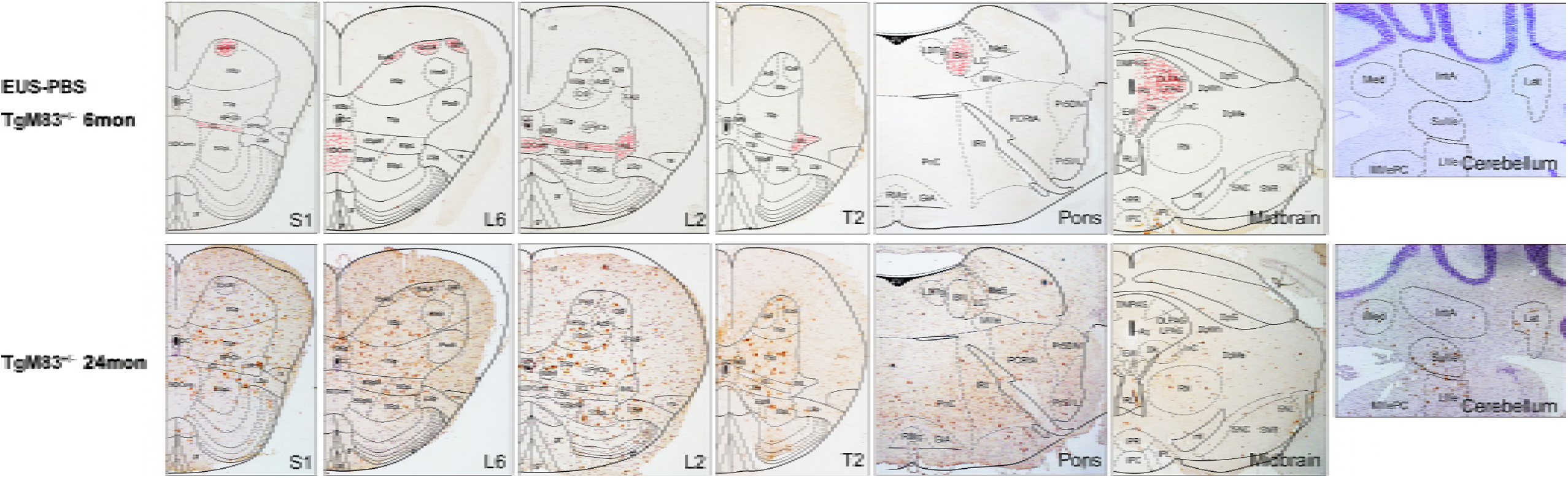
Representative immunohistochemical results of different segments from EUS-PBS TgM83^+/−^ mice and spontaneously ill TgM83^+/−^ mice. Sections were stained with anti-phospho-α-Syn (Ser 129) antibody. Representative images displayed the distribution of pα-Syn aggregates in S1, L6, L2, T2, pons, midbrain, and cerebellum. The locations, indicated by red dots in the upper panel, represent nuclei where pα-Syn aggregates were only observed in diseased EUS-α-Syn PFFs TgM83^+/−^ mice, but not in spontaneously ill TgM83^+/−^ mice.

**Figure 7-figure supplement 1.**
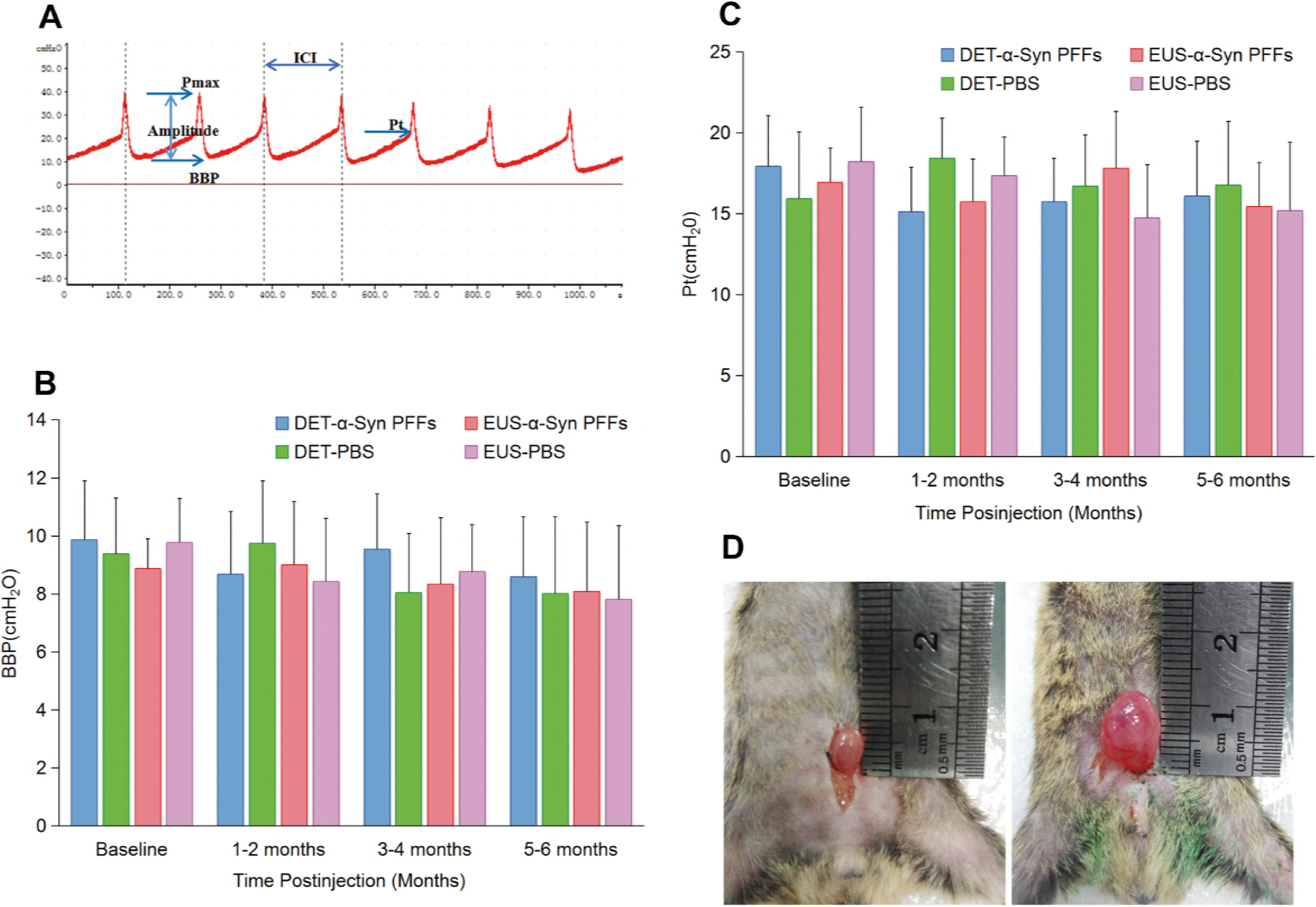
Urinary function analysis. (**A**) Representative cystometric curve of normal C57BL/6 mice. (**B** and **C**) Summary bar graphs from urodynamic evaluation about BBP (**B**) and Pt (**C**) for EUS and DET TgM83^+/−^ mice. EUS-α-Syn PFFs TgM83^+/−^ mice n = 18, DET-α-Syn PFFs TgM83^+/−^ mice n = 16, EUS-PBS TgM83^+/−^ mice n = 22, DET-PBS TgM83^+/−^ mice n = 20. Data are the means ± SD. Statistics was performed employing the Student’s t test and Mann-Whitney test. (**D**) Bladder size of EUS-α-Syn PFFs (right) and EUS-PBS (left) TgM83^+/−^ mice at 6-month post-injection.

**Figure 8-figure supplement 1.**
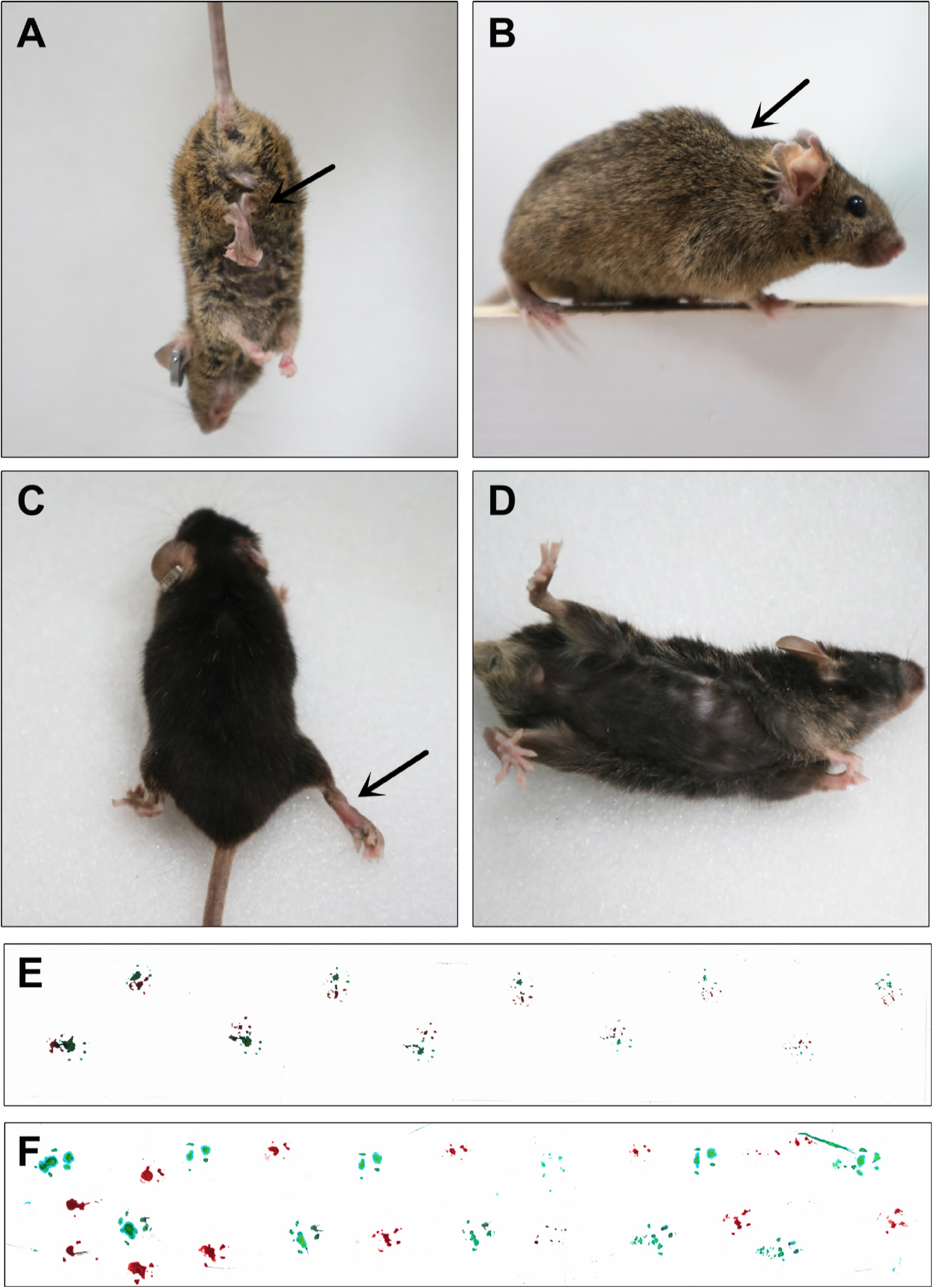
Motor and postural abnormalities of EUS-α-Syn PFFs TgM83^+/−^ mice at 5-month postinjection. (**A**) Hindlimb clasping (arrow). (**B**) Truncal dystonia (arrow). (**C**) Hindlimb dystonia (arrow). (**D**) Impaired postural adjustments. (**E** and **F**) Representative footprints of EUS-PBS TgM83^+/−^ mice (**E**) and diseased EUS-α-Syn PFFs TgM83^+/−^ mice (**F**).

**Figure 8-figure supplement 2.**
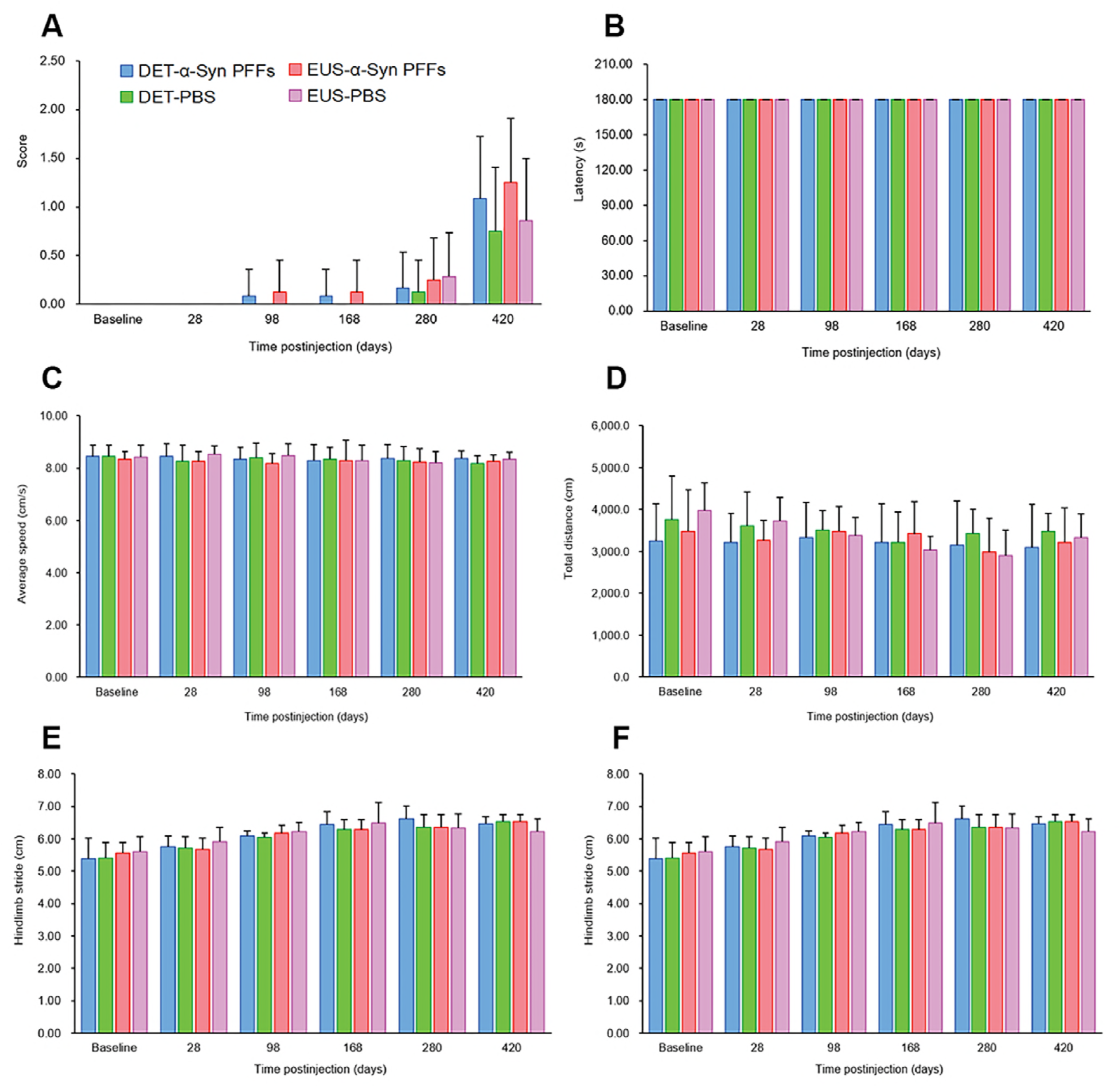
Behavioral analysis of C57BL/6 mice. (**A**) The mean score of MBS. (**B**) Latency to fall from the rotarod. (**C** and **D**) Average speed (**C**) and total distance (**D**) traveled during 15 minutes in the open field test. (**E** and **F**) Footprint analysis of the hindlimb stride length (**E**) and the hind-base width (**F**). EUS-α-Syn PFFs C57BL/6 mice n = 8, DET-α-Syn PFFs C57BL/6 mice n = 12, EUS-PBS C57BL/6 mice n = 7, DET-PBS C57BL/6 mice n = 8. Data are the mean ± SD. Statistical analysis was performed by using the Student’s t test and Mann-Whitney test, n.s., non-significant.

## Supplementary Materials

### Urodynamic examination (UE) and external anal sphincter electromyography (EAS EMG) of patients

All human subjects underwent clinical and electrophysiological evaluation, including UE and EAS EMG, as Yamamoto et al. previously described (Yamamoto et al., 2014), (Yamamoto et al., 2005).

### Surgery and retrograde tracing

To retrogradely label micturition reflex pathways, C57BL/6 and TgM83^+/−^ mice were anesthetized with isoflurane inhalation (S. Prusiner et al., 2015), and 15 µl Fluoro-Gold (FG) (Fluorochrome, LLC, Denver, CO) was injected slowly into EUS or DET. Following the injections, the skin was sutured. After 14 days, mice were perfused as described before (Bacskai, Rusznak, Paxinos, & Watson, 2014). The EUS or DET, pelvic ganglia, spinal cord and brain were removed and postfixed at 4 °C in a 30% sucrose solution containing 4% paraformaldehyde for at least 2 days. Serial transverse sections were cut at 20 µm using a freezing microtome (Leica CM1860 UV, Leica, Nussloch, Germany). Consecutive sections were collected, mounted, and cover-slipped with glycerol. Slides were then examined using an Olympus IX51 microscope equipped with epifluorescence.

### Transmission electron microscopy (TEM) imaging

The nature of the fibrillar α-Syn forms was assessed using Jeol 1400 (Jeol Ltd. Tokyo, Japan) TEM. First, a drop of fibrillar solution was transferred onto a carbon-coated 200-mesh grid and then negatively stained with 1% uranyl acetate. The images were recorded with Gatan Orius CCD camera (Gatan, Pleasanton, CA).

### Immunohistochemical and double immunofluorescence staining

Immunohistochemistry and double-labeling immunofluorescence analysis were conducted as previously described by Luk et al. (Luk, Kehm, Zhang, et al., 2012) using the following antibodies: phospho-α-synuclein (Ser 129) (mouse, Millipore, 1:600 or rabbit, Abcam, 1:400), α-synuclein filament (MJFR14) (rabbit, Abcam, 1:500), Ubiquitin (rabbit, Cell Signaling Technology, 1:400), Iba-1 (rabbit, Wako, 1:400), Myelin basic protein (rabbit, Abcam, 1:900), neurofilament heavy polypeptide (mouse, Abcam, 1:400), anti-tyrosine hydroxylase (rabbit, Abcam, 1:400), Rhodamine Red™-X (RRX) AffiniPure donkey anti-mouse IgG (H+L) (donkey, Jackson ImmunoResearch, 1:400), Cy™2 AffiniPure donkey anti-rabbit IgG (H+L) (donkey, Jackson ImmunoResearch, 1:400). Cell nuclei were stained using Hoechst33258 (1:1000, Solarbio). Slides were coverslipped with glycerol. Digital images were captured using Olympus IX51 microscope mounted with DP71 Olympus digital camera. Photoshop CS6 (Adobe Systems) was used to assemble montages.

### Western blotting analysis

After mice were anesthetized and decapitated, the spinal cord and several brain regions such as PAG, RN, pons, and cerebellum were separated on a cold stage. The isolated mice tissues were then stored in liquid nitrogen for further treatment. The transferred polyvinylidene fluoride (PVDF) membranes for the Western blotting as Kohl et al. previously described (Kohl et al., 2016) were incubated with following primary antibodies: phospho-α-synuclein (Ser 129) (mouse, Millipore, 1:600 or rabbit, Abcam, 1:800), α-synuclein filament (MJFR14) (rabbit, Abcam, 1:500), aggregated α-synuclein (5G4) (mouse, Millipore, 1:500). Forty-eight hours later, the membranes were washed in TBST (TBS with 0.1% Tween-20) and incubated with HRP-conjugated goat anti-mouse or goat anti-rabbit secondary antibodies for 2 hours at room temperature and visualized with enhanced chemiluminescence (Thermo Fisher Scientific). Proteins’ densities on the blots were normalized against those of GAPDH. All immunoreactive bands from Western blotting analysis were quantified by pixel intensity using FluorChem 8900 software (Alpha Innotech, San Leandro, CA, USA).

### Cystometry surgery

All animals were subjected to cystometric experiment to evaluate their urinary function before injection and at corresponding time points of post-injection following the methods reported previously (Boudes et al., 2013; Fandel et al., 2016; Girard et al., 2012; Silva et al., 2015). The animal was put supine and the bladder was exposed via a lower midline abdominal incision under isoflurane anesthesia. A polyethylene catheter-50 (Clay-Adams, Parsippany, New Jersey, USA) was implanted into the apical bladder dome and secured in place with a 6/0 purse-string sutures (Ethicon, Noderstedt, Germany). We flushed the catheter with saline to ensure no leakage and then threaded it from neck to the lower back through the subcutaneous tunnel anchored to the neck skin, finally closed the abdominal wall and skin. Through a three-way tap, the bladder catheter was connected to an infusion pump (B. Braun Sharing Expertise, Germany) and a pressure transducer (AD Instruments, Castle Hill, New South Wales, Australia) coupled to a computerized BL-420S data acquisition and analysis system (Techman Soft, Chengdu, China) which amplified and recorded intravesical pressure from the pressure transducer. We applied a heating lamp and room-temperature saline to maintain the body temperature of mice. Bladders were given a continuous infusion of 0.9% NaCl at a constant rate (20 µl/min) and after an equilibration period of 20-30 minutes, the intravesical pressure was recorded and voiding events were observed and noted for 30 minutes.

### Behavioral test

To evaluate α-Syn PFFs-induced behavioral deficits, mice were assessed by the following tests. TgM83^+/−^ and C57BL/6 mice were tested every 7 days starting from the second month’s post-injection. Blinded experiments were performed to treatment group for all behavioral tests.

### The motor behavioral scale (MBS)

MBS was used as Fernagut et al. previously reported (Fernagut et al., 2002). Higher score indicated higher disability and the maximum total score was 10. The total score was determined and used for the statistical analysis.

### Rotarod test

Motor coordination was assessed following the method previously reported by Duclot et al. (Duclot et al., 2012) with modifications. In brief, a rotating rod (Rotarod YLS-4C; YiYan Science and Technology Development Co., Ltd. Shandong, China) was used. At each time point, mice were placed on the rod rotating at 30 rpm. The latency to fall off the rotarod within the maximum time (180 seconds) was recorded, if a mouse stayed on the rod until the end of the 3 minutes, a time of 180 seconds was recorded. Mice received three trials per day with a 15-minute inter-trial interval. The mean latency to fall off the rotarod was statistically analyzed.

### Open field test

To assess general activity, locomotion, and anxiety of the mice, the open field test system (Wuhan YiHong Sci. & Tech.Co., Ltd) was applied. Mice were placed in the center of the open field (37.5 × 37.5 × 34.8 cm) and tested for 15 minutes at the same time of the day (6:00 p.m. to 9:00 p.m.). Activity was analyzed by the Anilab software version 5.10, registered version (Anilab Software & Instruments Co., Ltd., China). At the end of testing, the arena was cleaned with 75% alcohol to remove olfactory cues. The tests were performed in a dark room that was isolated from external noises and light during the test period. Total distance (cm), average speed (cm/s), and zone crossing were statistically analyzed.

### Footprint test

The footprint test was performed to examine the gait of the mice. Paws of the mice were painted with water-soluble non-toxic paint of different colors (fore-paws in red and hind-paws in green). The animals were then allowed to walk along a restricted cardboard tunnel (50 cm long, 5 cm wide, 10 cm high) into an enclosed box and a sheet of white paper (42 cm long, 4.5 cm wide) was placed on the floor of the tunnel, and one set of footprints was collected for each animal. Three steps from the middle portion of each run were measured for four parameters (cm): (1) stride length (front and hind legs). (2) The front-and hind-base width. The mean of each set of values was statistically analyzed (Stefanova et al., 2005).

### Beam walking test

Balance and bradykinesia were assessed with the method described before with modifications (Schafferer et al., 2016). The beams consisted of two different types of wood (each measuring 80 cm long, one was 1.6 cm, and the other 0.9 cm wide) placed horizontally 50 cm above the floor, respectively. Two daily sessions of three trials were performed using the 1.6 cm width large beam during training. Mice were then tested using the 0.9 cm width beam. Mice were allowed to perform in three consecutive trials. The time for traversing 50 cm as well as the number of sideslip errors were recorded on each trial. The average traverse duration and average number of sideslip errors of the three trials were statistically analyzed.

### Pole test

The pole test was performed to assess motor coordination and balance. A vertical gauze-taped pole (1 cm diameter, 50 cm height) with a small cork ball (3 cm diameter) at the top was applied. Mice were placed with their head upward right below the ball. The time taken to turn completely downward (T turn) and total time taken to reach the base of the pole with four paws (T total) were recorded. The maximum cutoff of total time to stop this test was 120 seconds. This test was performed three times for each mouse, while the average time was statistically analyzed (Zhou et al., 2016).

